# Spatially-resolved single-cell atlas of ascidian endostyle provides insights into the origin of vertebrate pharyngeal organs

**DOI:** 10.1101/2023.05.03.539335

**Authors:** An Jiang, Kai Han, Jiankai Wei, Xiaoshan Su, Rui Wang, Wei Zhang, Xiawei Liu, Jinghan Qiao, Penghui Liu, Qun Liu, Jin Zhang, Nannan Zhang, Yonghang Ge, Yuan Zhuang, Haiyan Yu, Shi Wang, Kai Chen, Xun Xu, Huanming Yang, Guangyi Fan, Bo Dong

## Abstract

The pharynx is an endoderm innovation in deuterostome ancestors, the vertebrate descendent structure of which is a pharyngeal developmental organizer involving multi-germ layer and organ derivatives. However, the evolutionary origination of complicated pharynx organs in vertebrates is still largely unknown. Endostyle, a transitional pharyngeal organ exclusively in basal chordates provides an opportunity to reveal the origin of pharyngeal organs. Here, utilizing cutting-edged Stereo-seq and single-cell RNA-seq, we constructed the first spatially-resolved single-cell atlas in the endostyle of urochordate ascidian *Styela clava*, where the spatial location of Stereo-seq and high capture efficiency of single-cell RNA-seq complement each other and identified 23 highly differentiated cell types. We identified a previously overlooked hemolymphoid region (HLR), which harbors immune and blood cell clusters with enriched stemness capacities, illuminating a mixed rudiment and stem-cell niches for the blood and lymphoid system. More excitingly, we discovered a mechanical-sensitive hair cell candidate in zone 3 homologous to vertebrate acoustico-lateralis system, which was supported by the expression of *in situ* hybridization-verified inner ear-specific markers, including *PTPRQ*, *USH2A*, *WHRN*, and *ADGRV1*, ultracellular structure evidence and cross-species comparison. These results thoroughly renewed the comprehension of the basal-chordate pharynx and provides expressional evidence for multiplexed pharyngeal organ evolution.

## Main

The development of pharyngeal organs, including gill slits and apparatus, is the most prominent morphological innovation for deuterostome ancestors^1–3^, which was described in stem-echinoderms^4^ and stem-deuterostomes^5^. The most primitive form of the pharynx in extant deuterostome has been described in hemichordates as simple pharyngeal endodermal out pockets from foregut^6^. While the pharyngeal endoderm in vertebrates shared a homologous developmental origin and regulatory network with that in the primitive form^2, 7^, which is an organizer for the vertebrate pharyngeal framework^8, 9^. The complexity of the pharynx in vertebrates arose with the incorporation of cranial paraxial mesoderm and neural crest-derived mesenchyme, which gives rise to series of appendix organs within pharyngeal arches. Besides the thyroid developed from the pharyngeal endoderm, multiple specialized appendices were derived from the pharynx primordium, like the parathyroid, the thymus, and the auditory organ^10^.

The endostyle is a specialized pharyngeal organ possessed exclusively by non-vertebrate chordates and larvae of ammocoetes, evolutionary transitional hierarchies between vertebrates and non-chordate deuterostomes. It takes the form of a groove-like channel that longitudinally extends through the ventral body wall, both lateral of which is composed of regional ciliated epithelial cells specialized in mucus secretion and histological support^11^, facilitating the process of filter-feeding^12^. As an endoderm-derived organ^13^, regional-specific markers labeling showed the tissue expressional specificity and pattern formation features in the endostyle^14–16^. The endostyle has been recognized as a thyroid equivalent organ because of cellular components capable of iodine concentrating^17, 18^ and thyroid hormone synthesis^14, 15, 19^. Recent studies have shown that the endostyle may have diverse functions and possible evolutionary prototypes for organs in vertebrates. In colony ascidian, the sinus region has been proved with hematopoietic functions^20^ and stem-cell harbor features^21^. The peptidergic neuron has been reported to exist in the endostyle sinus regions^22^. Additionally, the endostyle and its surrounding sinus have been reported with immune functions, evidenced by immunohistochemical labeling under pathogen-origin bio-chemical compound threats^23, 24^. Overall, these reports highlight the potential that candidate cellular compositions of multiple functions may exist, implying the endostyle as a multiplexed organ with functions of a pharynx prototype in basal chordates.

In complex organ evolutionary studies, cell type definition and comparison are the cornerstone for functional comparison^25^. With the aid of advancing single-cell transcriptomics, researchers are now capable of defining the cellular composition of an organ at the single-cell resolution^26–28^. However, for organs of specialized tissue arrangement patterns, like the endostyle, it is necessary to detect the transcriptome in a location-addressable manner. To comprehensively investigate the cellular composition and evolutionary status of the endostyle, we utilized a cutting-edge spatial transcriptomic technic, Stereo-seq^29^, and high throughput single-cell transcriptome method^30^, 10x genomics to construct a location-addressable cellular transcriptional profile for the endostyle transverse sections in a typical urochordate ascidian *S. clava*, whose genome and transcription factors have been well characterized^31, 32^. In total, we constructed a spatially-resolved atlas of the endostyle and defined 23 cell clusters by integrating three bio-replicons for single-cell RNA-seq and six Stereo-seq sections. We also performed electron microscopy observation, verification of marker gene expression, and the comprehensive cross-species analysis of crucial cell types. Based on these data, we identified a new hemolymphoid-holding region as a part of endostyle and discovered novel functional cell types in the endostyle, such as a cell population in zone 3 with representative markers of mechanical-sensitive hair-cell architecture and neuron transduction, which is probable to be the acoustico-lateralis system homolog. Our research comprehensively uncovered the cellular composition of an evolutionary transitional pharyngeal organ, the endostyle, and largely enriched our knowledge of cellular population foundation deriving complex advance organs in vertebrates.

## Results

### Pharynx-related diverse cellular compositions in ascidian endostyle

The pharynx exists in hemichordates, basal-chordates, and vertebrates in extant deuterostomes (Fig.1a). The hemichordates possess the simplest pharyngeal structure, gill slits originating from endoderm out pockets. Whereas in vertebrates, besides feeding and respiration tunnel, multiple derivative organs with versatile functions developed from the pharyngeal arch, including the auditory organ, thyroid, tonsil, and thymus. In the evolutionary transitional category between invertebrates and vertebrates, the endostyle, a distinct pharyngeal organ, constructs the pharynx together with gills.

The transverse section of the endostyle is composed of diversified regional components. A simple bilateral symmetric groove is half-embraced by nine ciliated epithelial on lateral sides, including the surrounding sinus region adjacent to the ventral blood vessel (VBV) (Fig. 1b). Longitudinal levels are largely homogenous except for the difference of a few *Hox* genes^33^. To uncover cellular components and potential vertebrate homologs in the endostyle, we constructed a spatially-resolved atlas at single-cell resolution (Fig. 1b and Extended Data Fig.1a). The single-cell RNA-seq dataset, containing 10,017 valid cells, was processed with Seurat workflow and cell compositions were preliminarily defined (Extended Data Fig. 1b). Multiple cell types including immune cells, secretory epithelial cells, and blood cells were discovered with cluster-specific markers (Extended Data Fig. 1c and Supplementary Table 1). In Stereo-seq, the expression profile with spatial location detected by DNB (DNA nano-ball) was aligned against the optical image captured by staining with ssDNA dye in library construction (Extended Data Fig. 1d). Then, two cell segregation strategies, including squared bin division and cell segregation, were applied to the dense tissue region and the sparse tissue region respectively in dividing DNB spots into cell units (Extended Data Fig. 1e). Cell unit boundaries segregating DNB spots on silicon chips were constructed as the atlas backbone (Extended Data Fig. 1f). After cell segregation, 18,371 cell units were obtained in six Stereo-seq sections and cell type annotation were preliminary conducted (Extended Data Fig. 1g and Supplementary Table 2).

**Fig 1:**
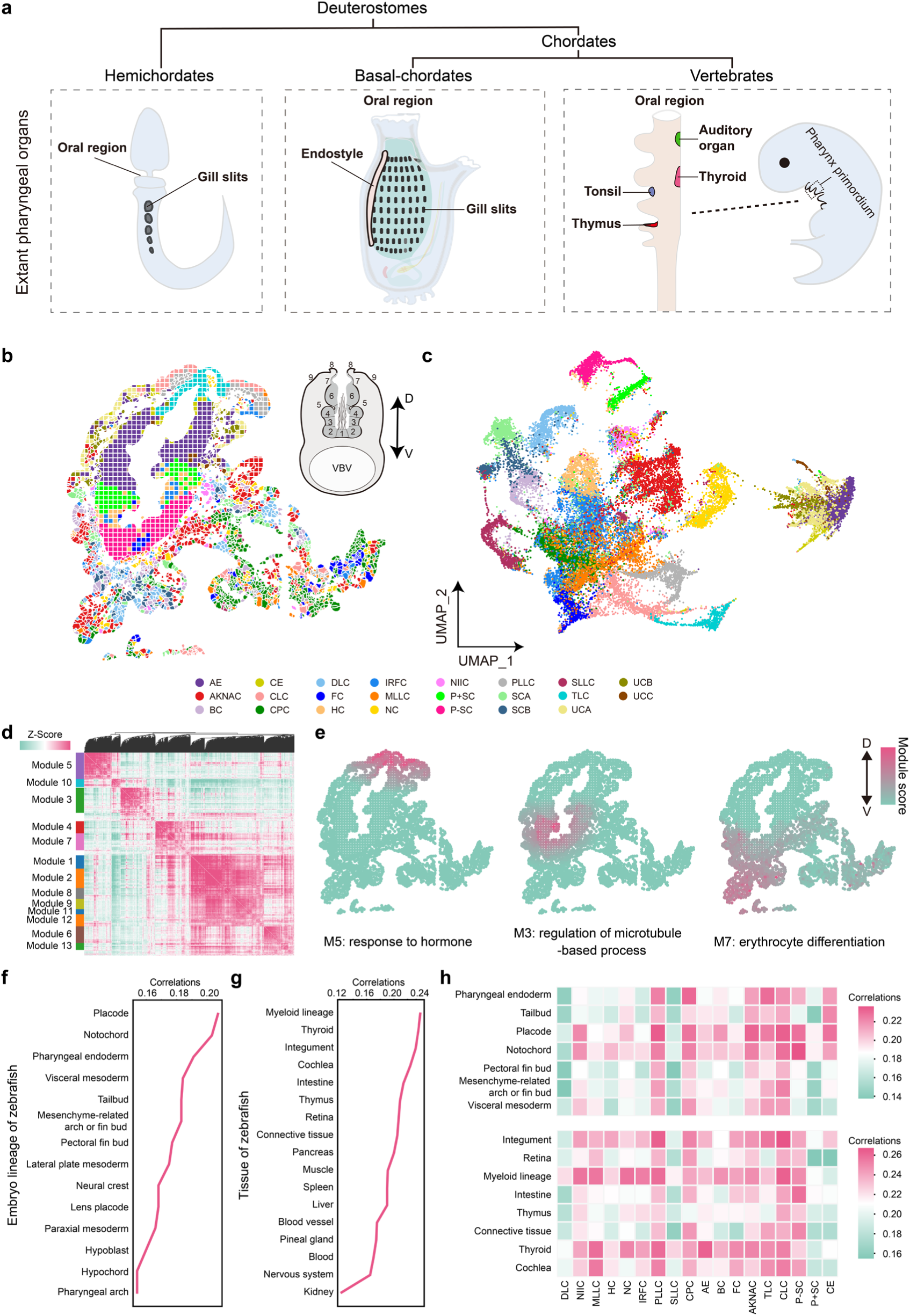
Cell atlas construction for endostyle in adult basal-chordate, *Styela clava*. **a**, Extant pharyngeal organs and apparatus in three deuterostomes categories, hemichordates, basal-chordates, and vertebrates. Left, schematic of pharynx organ, gill slits in hemichordates. Middle, schematic of pharynx organ in basal-chordates, represented by ascidian, including gill slits (on pharyngeal sac) and the endostyle. Right, schematic of the pharynx primordium in vertebrates. Pharynx organs (tonsil, thymus, auditory organ, and thyroid) have primordium in the pharynx arch. **b,** Spatial atlas shows the distribution of cell units on the transverse section colored with cell annotation. AE: Antimicrobial epithelium, AKNAC: AKNA+ cell, BC: Blood cell, CE: Connective epithelium, CLC: Ciliated cell, CPC: Complement cell, DLC: Dendritic-like cell, FC: Flagellate cell, HC: Hair cell, IRFC: IRF+ cell, MLLC: M3K1+ lymphoid like cell, NC: Neuroendocrine cell, NIIC: Neuro-immune interact cell, P+SC: PPIB+ secretory cell, P−DC: PPSB− secretory cell, PLLC: PA21B+ lymphoid like cell, SCA: Styelin cell A, SCB: Styelin cell B, SLLC: STYC+ lymphoid like cell, TLC: Thyroid like cell, UCA: Undefined cell A, UCB: Undefined cell B, UCC: Undefined cell C. Upper right schematic showing the transverse structure of the endostyle, consisting bilateral symmetric zone 1-9 with a ciliated groove in the center, surrounded by the sinus region. The subendostylar vessel exists on the ventral side. D: dorsal, V: ventral. **c,** UMAP of all 28,388 scRNA-seq and stereo-seq cell units, colored with cell annotation. **d,** Heatmap showing gene modules based on spatial autocorrelation. **e,** Spatial visualization of selected modules in the heatmap of gene modules. Three exhibited modules were labeled with the enriched function of genes. D: Dorsal, V: Ventral. **f and g,** Broken lines showing spearman correlations between the expression profile of the endostyle and embryo lineages (**f,**) or tissues (**g,**) of zebrafish (*Danio rerio*). **h,** Heatmap showing spearman correlations between the expression profile of cell clusters in the endostyle and zebrafish tissues.

Then, we comprehensively integrated the dataset from three single-cell RNA-seq bio-replicons and Stereo-seq from six sections of 10 μm thickness (Fig. 1c, Extended Data Fig. 2) and annotated cellular types *de novo* based on the cluster-specific expressed gene list (Extended Data Fig. 3a and Supplementary Table 3). Annotation was projected on spatial landscapes to construct the cell atlas (Fig. 1b). It’s clearly shown a bilaterally symmetric pattern with homogenous cell-type composition in dense tissue region (DTR) and scattered cell patterns in the surrounding region. As the immune and blood cell components in the surrounding sinus, we defined it as the hemolymphoid region (HLR) of the endostyle in this research. To evaluate the batch effect of two technical repeats, we first compared the cell numbers of cell clusters in the atlas. The result showed a general homogenous feature in cell number distribution among clusters (Extended Data Fig. 3b). For the UMAP plot separated by the technical batch, a well-corresponded pattern was seen in most cell clusters (Extended Data Fig. 3c). We also independently annotated cell type in single-cell RNA-seq data and Stereo-seq data respectively, and projected annotations onto the UMAP space. Most regions of the two technical batches showed direct cell type correspondence (Extended Data Fig. 3d).

**Fig 2:**
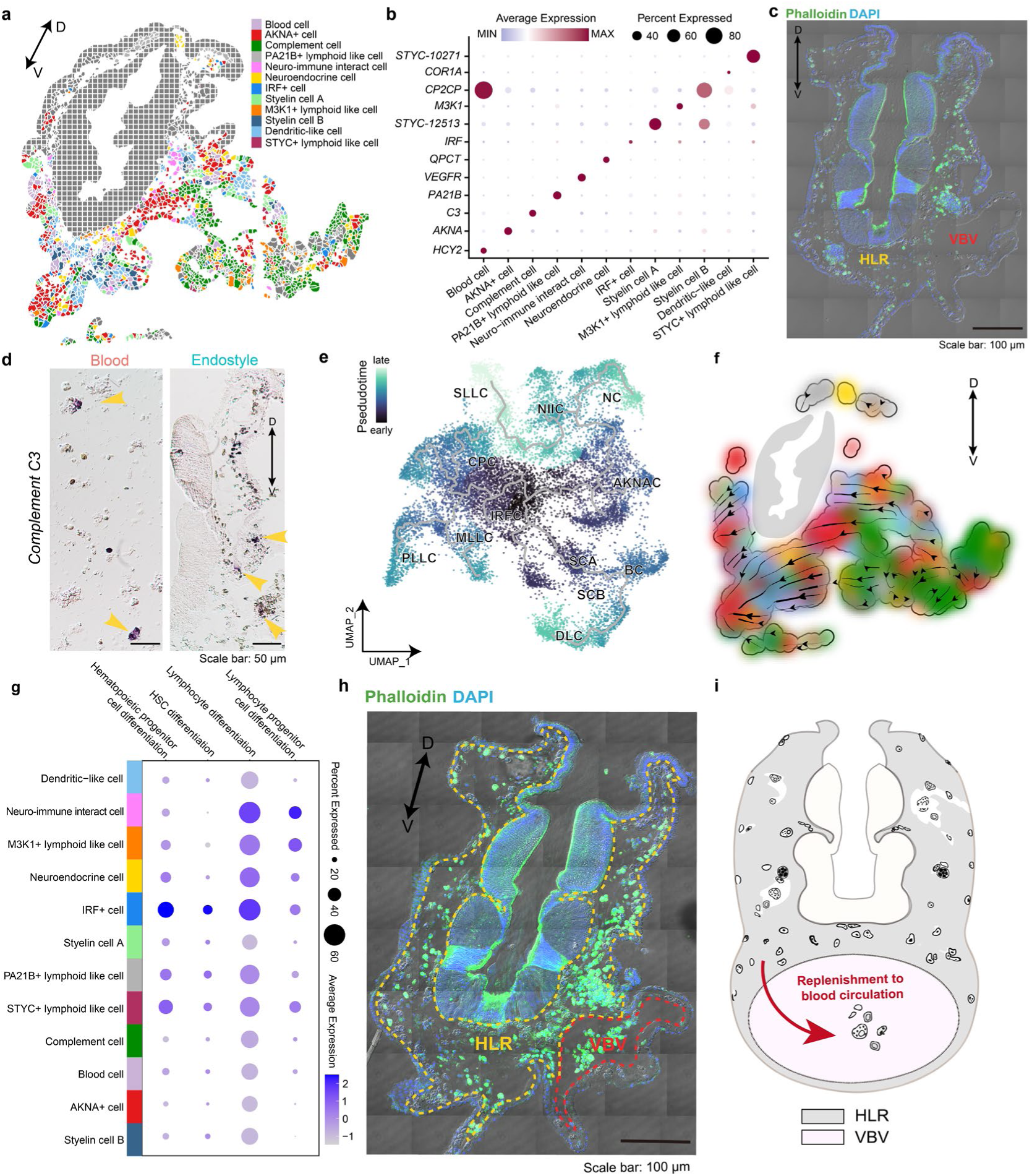
Hemolymphoid like region (HLR) provides a lymphoid-like niche for blood/immune cell lineages. **a**, Spatial visualization of cell clusters in HLR. D: dorsal, V: ventral. **b,** Bubble plot showing the expression of marker genes in indicated cell types. **c**, Structure and location of HLR with results of confocal microscopy. **d,** *In-situ* hybridization showing the expression pattern of *Complement C3*. Left, result on the peripheral blood cell. Right, result on the endostyle cryosection. D: dorsal, V: ventral. Yellow arrows point at signals. **e,** Pseudotime trajectory analysis depicting trajectories among cell clusters in the HLR based on the UMAP. Cells are colored with pseudotime value on the UMAP. **f,** RNA velocity streamline plots showing the predicted trajectory of cell clusters transition in the HLR of the endostyle. Cells are plotted based on the location in spatial atlas. D: dorsal, V: ventral. **g,** Bubble plot showing expression intensity of HLR cell clusters in exhibited gene categories. **h,** Confocal result showed scattered distributed cells in HLR. VBV, ventral blood vessel. **i,** The model for immune and blood system in HLR.

**Fig 3:**
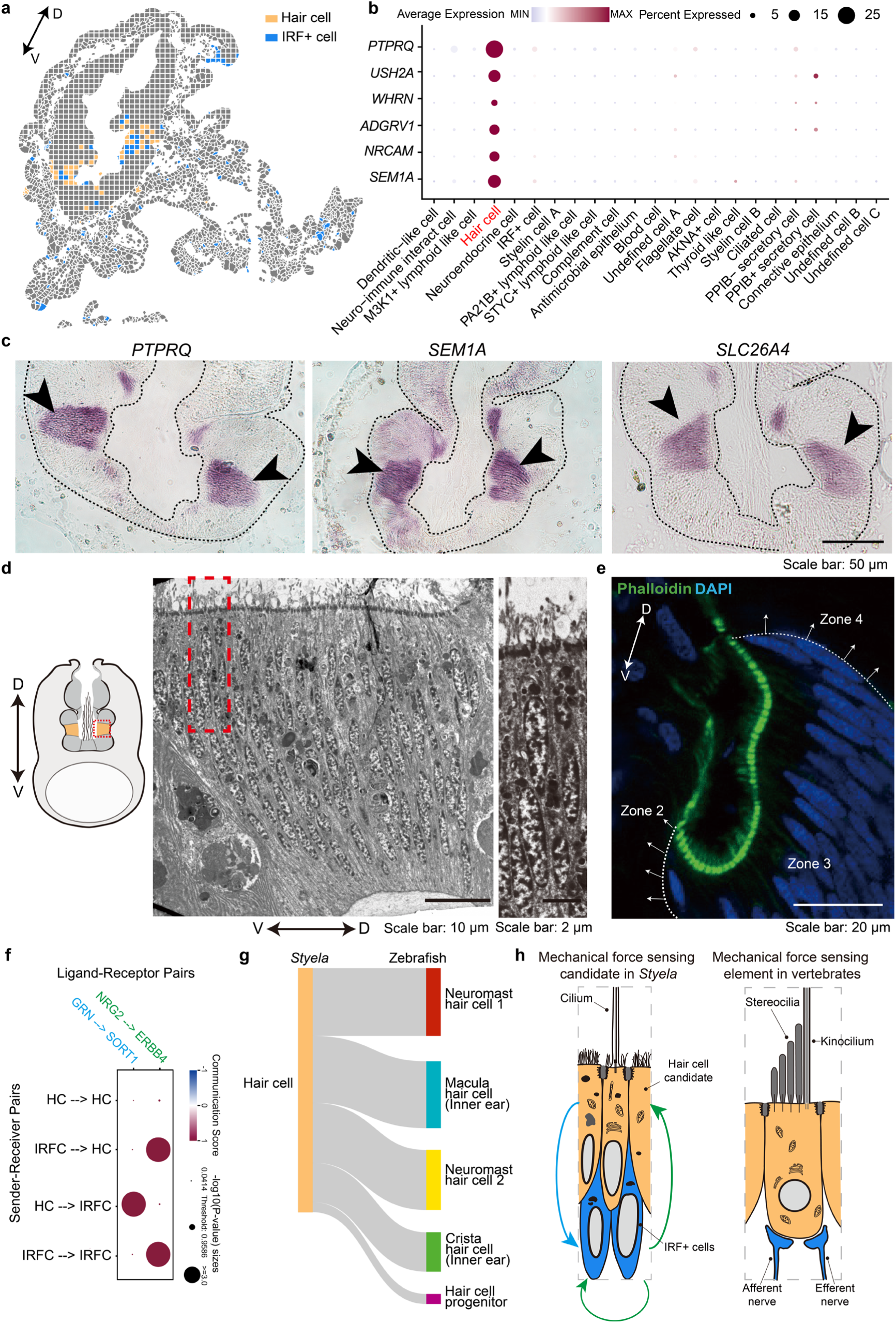
Mechanical-sensitive cell cluster defined in zone 3 as the acoustico-lateralis homolog candidate. **a**, Spatial visualization of hair cell cluster and IRF+ cell cluster. D: dorsal, V: ventral. **b,** Bubble plot showing the expression of marker genes in indicated cell types. **c,** *In-situ* hybridization verified the expression pattern of *PTPRQ*, *SEM1A* and *SLC26A4*. Black arrows point at signals. **d,** Left, schematic for the location of zone 3 in the endostyle transverse section (orange region). Middle, Detailed visualization of subcellular structure in the region of red rectangle in the left figure. Right: More detailed visualization of the red rectangle region in the middle figure. D: dorsal, V: ventral. **e,** Confocal microscopy visualized the zone 3 region on Phalloidin (green) and DAPI (blue) stained cryosection. Zone 2 to 3 and zone 3 to 4 tissue boundaries were drawn with white broken lines. D: dorsal, V: ventral. **f,** Bubble plot showing communication scores of ligand-receptor pairs between sender and receiver cell populations. HC: Hair cell, IRFC: IRF+ cell. **g.** Cross-species comparison between hair cell cluster and cell populations of the acoustic-lateralis system in zebrafish. **h,** Left, schematic for mechanical force sensing candidate in the endostyle, blue arrow and green arrow corresponding to ligand-receptor relationship, GRN to SORT1 and NRG2 to ERBB4, respectively. Right, schematic for mechanical force sensing element in vertebrates.

To describe potential functions from the expression level, we first selected high-variable genes in the integrated dataset (Extended Data Fig. 3e). GO enrichment analysis revealed highly variable functions in the endostyle, including neuron transduction (axon guidance and glial cell differentiation), immune process (adaptive immune system), muscle (muscle filament sliding), and pathways relating to microtubule cytoskeleton or supramolecular fiber organization (Extended Data Fig. 3f). Moreover, we identified 13 gene modules with significant spatial autocorrelation in the Stereo-seq dataset with Hotspot^34^ (Fig. 1d). Module scores representing the expression level of genes were projected on the spatial landscape. The functions of modules were defined with top GO enrichment terms. Heatmaps exhibited regional intensive functions, including response to hormone, regulation of the microtubule-based process, and erythrocyte differentiation in dorsal parts of DTR, ventral parts of DTR, and HLR, respectively, which is parallel with the function of hormone metabolism, the mucus-net for filter-feeding^35^, and surround blood region in HLR^20^ (Fig. 1e).

To comprehensively describe the potential homologous between the endostyle and vertebrate, we summarized and reanalyzed the single-cell dataset from zebrafish (*Danio rerio*)^36–38^. Expressional correlations between the endostyle and embryonic lineages/tissues from zebrafish were evaluated with the Spearman algorithm based on the average expression of 387 homologous transcription factor (TF) genes in *S. clava* and *D. rerio* (Supplementary Table 4). The results for embryo lineage of zebrafish showed complex germ-layer similarity for the endostyle, including placode, notochord, pharyngeal endoderm, and visceral mesoderm (Fig. 1f), most of which are key germ-layer components of the pharyngeal arch in development^39^. Moreover, the comparison to the tissue of zebrafish showed high similarity of the endostyle to myeloid lineage, thyroid organ, integument, and so on (Fig. 1g), hinting at potential similarity between the endostyle and specialized organs. The correlation between the endostyle and tissues from zebrafish was expanded to the cell type level (Fig. 1h).

### HLR is a homolog candidate for blood/immune organs in vertebrates

To fully identify the cellular composition of the HLR region in the spatially-resolved single-cell atlas, we first did cell selection based on cell type annotation and distribution (Extended Data Fig. 4a). We selected cell clusters dominated by cell units located in the sparse region of Stereo-seq results. All single-cell RNA-seq and Stereo-seq cell units in sparse regions of these clusters were included and visualized for further analysis (Fig. 2a). Clusters-specific markers were exhibited in supporting cell type annotations, most of the clusters were immune/blood functional related (Fig. 2b). Expression patterns of markers, *C3* and *HCY2*, were verified by *in situ* hybridization (ISH) (Extended Data Fig. 4b).

**Fig 4:**
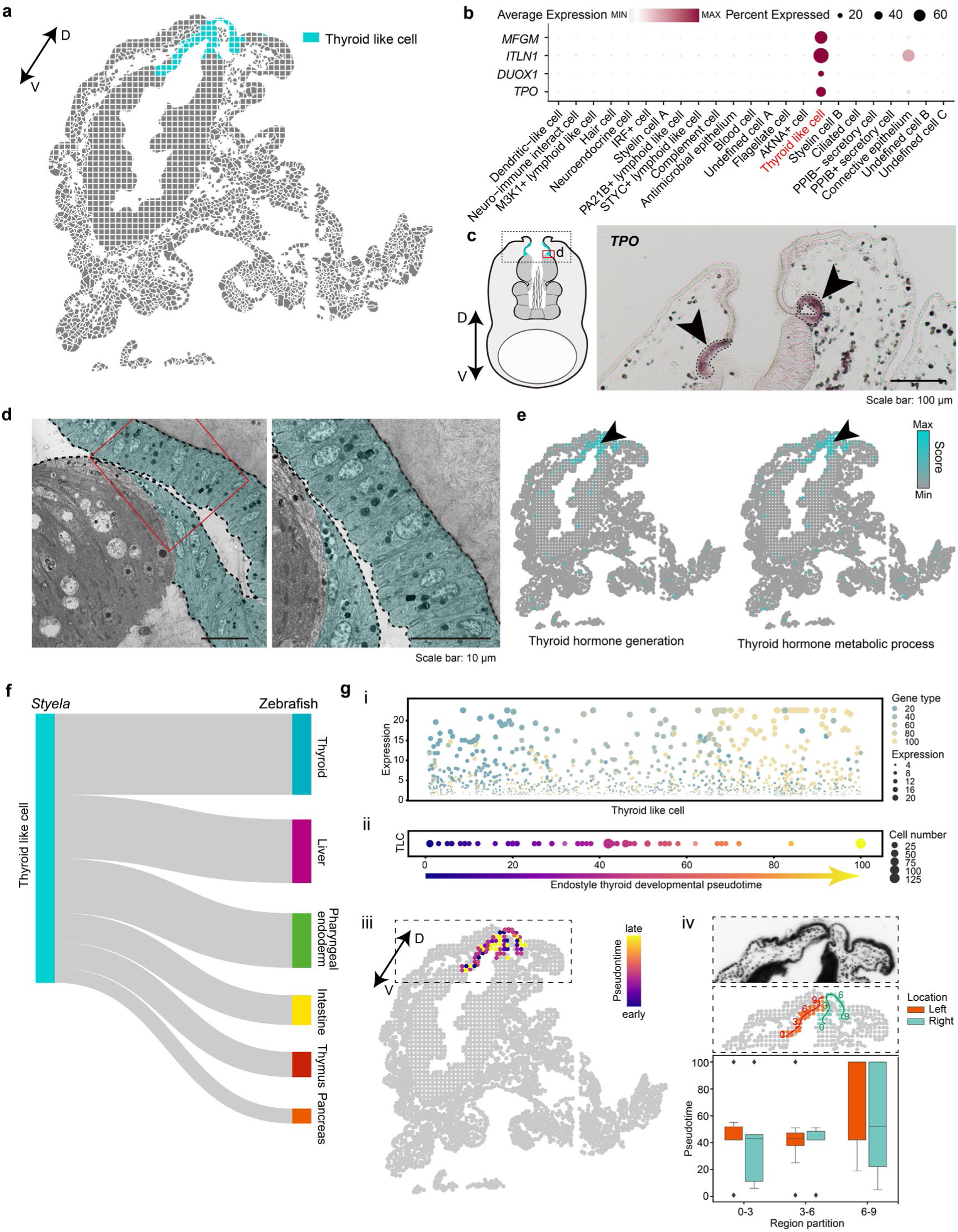
Homologous cell cluster to thyroid gland in the endostyle cell atlas. **a**, Spatial visualization of thyroid like cell (TLC) cluster. D: dorsal, V: ventral. **b,** Bubble plot showing the expression of marker genes for TLC, including *TPO*, *DUOX1*, *ITLN1*, and *MFGM* in indicated cell types. **c,** Left, schematic for the location of TLC in the endostyle transverse section. Right, in-situ hybridization showing the expression pattern of *TPO*. Black arrows pointing at signals. The field of view is the black rectangular region in left schematic. **d,** Left, TEM visualizes the ultracellular structure of cells in TLC cluster (masked with blue). Right: Detailed visualization of red rectangle in the left figure. **e,** Heatmap showing expression of indicated functional pathways in the spatial plot. **f,** Cross-species comparison between the TLC cluster in the endostyle and tissue/organs in zebrafish. **g, (i)** Expression of genes with trajectory information (Extended Data Figure 5d, 5e) in cells of TLC. Each bubble representing the expression level of certain gene in one cell. Bubbles are colored with the trajectory location of the gene, also the pseudotime position of the gene. The size of bubbles showing its expression level. Cells in TLC cluster were ranked on horizontal axis based on the trajectory type of their top expressed genes. **(ii)** Distribution of cells in TLC on the pseudotime trajectory defined by the trajectory type of their top expressed genes. Bubble size indicates the cell number in a certain pseudotime status. **(iii)** Trajectory statues visualization on spatial atlas, the color corresponding to the pseudotime value. **(iv)** Upper, the ssDNA image for rectangular region in (iii), Middle, track line showing the simple epithelial extending from the ventral region to dorsal region closing to zone 8, which is divided into nine segments evenly. The left and right zone 7 were label with different colors respectively. Lower, barplot showing the statistics of pseudotime values for cells on nine segments.

To uncover the morphological features of HLR, we conducted the confocal microscopy and transmission electron microscopy (TEM) morphological observation on the region. The result showed three tissue components, HLR, and VBV on the transverse section (Fig. 2c). The HLR was largely filled with a highly developed extracellular matrix (ECM), which formed caved-like structures. Large number of cells were scattered in the ECM space (Extended Data Fig. 4c). We systematically compared the scattered distributed cells in HLR to the blood system in ascidian^40^. Most types of blood cells were identified according to cell morphology in all nine cell types in the blood system of *Ciona robusta* (Extended Data Fig. 4d).

To verify the potential divergence of the HLR and peripheral blood system, we utilized ISH to detect the expression of marker gene *Complement C3*, for the complement cell cluster in HLR scattered cells and the peripheral blood cell, respectively (Fig. 2d). Statistical result showed that the ratio of C3+ cells population in HLR were significantly larger than that in peripheral blood tissue, which proved the enrichment of immune-labeled cells in HLR compared to peripheral blood, suggesting the involvement of intensive immune activity in HLR (Extended Data Fig. 4e).

Next, a pseudotime analysis was conducted on cell clusters in HLR. Cells from the single-cell dataset with the annotation of ‘stem cell’ were assigned as the root of pseudotime trajectory, which are strongly capable of proliferating (Extended Data Fig. 1c). The result showed the complement cell and IRF+ cell cluster were of the lowest pseudotime value, suggesting their primary differential status. While clusters of terminal developmental states were supported by high pseudotime values, including dendritic like cell, blood cell, and AKNA+ cell clusters (Fig. 2e, Extended Data Fig. 4f). RNA velocity analysis with dynamo also showed a similar evolving trend with results from Monocle 3. The general trend of streamlines showed origination from the complement cell cluster and IRF+ cell cluster (Extended Data Fig. 4g). Then, streamlines of RNA velocity were projected onto the spatial landscape, which showed a general trend of evolving from the ventral part of HLR to dorsal regions (Fig. 2f).

To probe the potential stemness of blood/immune lineages in the HLR, we conducted expression visualization on stemness marker gene categories. The expression of gene categories in the HLR clusters was visualized by expression bubble plot (Fig. 2g). The IRF+ cell has an intensive expression of genes in HSC differentiation and hematopoietic progenitor cell differentiation categories. Cell populations including M3K1+ lymphoid like cell, STYC+ lymphoid like cell, and PA21B+ lymphoid like cell have enriched expression in genes of the lymphocyte differentiation category. These results revealed cell clusters with lower pseudotime values or immature status, have higher stem capacity. Meanwhile, it also showed a developmental trend that primitive cells were prone to the ventral region of the HLR while cells in the dorsal part were prone to the terminal developmental trajectory.

These new insights have revealed that HLR is a hub for the generation and maintenance of blood and immune cells. The HLR is believed to be a mixture of primordial hemolymphoid system that shares homology with the blood and immune systems of more advanced species. Therefore, we consider the HLR to be a key region having stem cell center which give rising to functional cells of both the immune and blood systems, might be critical for replenishing blood circulation (Fig. 2h and 2i).

### The hair cell cluster in zone 3 is a homolog candidate for acoustico-lateralis system

Hair cells were identified in zone 3 of DTR together with the IRF+ cells (Fig. 3a). The hair cell cluster showed specific expression of markers including *PTPRQ*, *USH2A*, *WHRN*, and *ADGRV1*, which are key elements in maintaining functionally active hair bundle and stereocilia in hair cells of the vertebrate^41–45^ (Fig. 3b, Extended Data Fig. 5a). Meanwhile, specific expressions of neural functional markers, *NRCAM*, *SEM1A*, and *SLC26A4*, were also distinctively detected. *NRCAM* is of vital importance in the development of spiral ganglion neurites^46^. *SEM1A* plays a role in growth cone guidance^47^. *SLC26A4* is an electroneutral sodium-independent transported of chloride and iodide relating to deafness and Pendred syndrome^48^. The expression pattern of *PTPRQ*, *SEM1A*, and *SLC26A4* were verified with ISH, showing distinct signals in zone 3 (Fig. 3c, Extended Data Fig. 5b). For the coexisted cell population, IRF+ cell in zone 3, distinct markers compared to the hair cell cluster were *GSN* (Gelsolin), *ACTN2*, *IF* (Intermediated Filament), *PLS3*, and *AXNA7*, many of which are important cytoskeletal components, which may correlate with morphological construction and mechanical supporting (Extended Data Fig. 5c).

**Fig 5:**
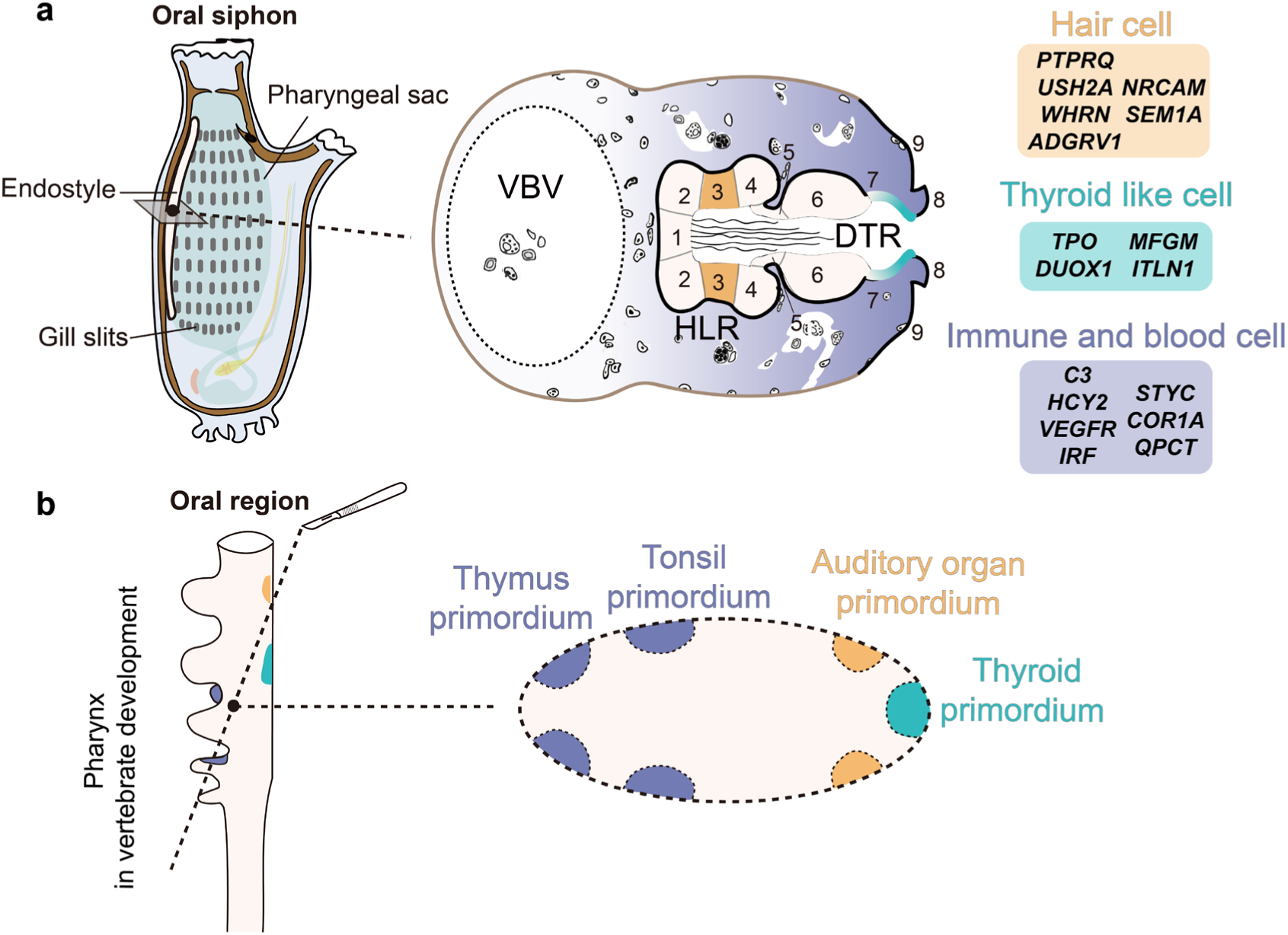
Comparison of the pharynx organ between the endostyle in ascidian and developing pharynx in vertebrates. **a**, Left, schematic showing the location of the endostyle in adult ascidian. Right, the transverse section of the endostyle showing the diverse cellular composition of the endostyle, including the immune and blood cell, the hair cell, and the thyroid like cell, which corresponding main cell populations for the HLR, zone 3, and zone 7 in the schematic. The gradient color in three regions showing their diverse developmental status within tissue. Distinct marker genes were label besides. VBV, Ventral blood vessel. HLR, hemolymphoid region. DTR, dense tissue region. **b,** Left, schematic showing the morphology illustration for developing pharynx in vertebrates. Right, the diagonal plane of the developing pharynx showing multiple organ primordium, including thyroid, auditory organ, tonsil, and thymus.

To describe the morphological features and relationship of hair cell and IRF+ cell populations, we conducted TEM and confocal microscope visualization. TEM results showed that the cells in zone 3 were in narrow shuttle shapes interleaving tightly with each other (Fig. 3d, Extended Data Fig. 5d-5e). Long F-actin cilia were clearly distributed on the apical surface with dense narrow cell nuclear arranging below (Fig. 3e). The hair cell population with distinct apical cilia connected tightly with basal cells, which were thought to be IRF+ cell. The relationship between two cell population were clearly visualized with ligand-receptor analysis showing interaction from NRG2 to ERBB4 and GRN to SORT1, both of which are important cell-cell interactions in neural function^49, 50^ (Fig. 3f and Extended Data Fig. 5f). To investigate the potential evolutionary relationship of the hair cell candidate in zone 3 with organs/tissue in vertebrates, we conducted a cross-species comparison analysis. The result showed that the hair cell cluster in zone 3 has high expression correlations with hair cell populations and hair cell progenitor of the acoustico-lateralis system in zebrafish (Fig. 3g).

Based on expressional and morphological results, we promote a mechanical force sensing model, which is presumed to be the most ancient homolog to the vertebrate inner ear and aquatic acoustic-lateralis system^51^. Contrasting with mechanical force sensing elements composed of hair cells and synapse on basal membrane in vertebrates, hair cell and IRF+ cell populations in *Styela* endostyle constructed two tightly interleaved layers from apical ciliated surface to basal supporting cells with tight junction-dependent transmission capacity of nerve electrical signals in the absence of synapse structure (Fig. 3h).

### Coexistence of different maturation status of thyroid like cell in the thyroid-equivalent element in zone 7

A thyroid like cell (TLC) cluster was defined in zone 7 of the endostyle in the spatial atlas (Fig. 4a). The cell cluster detected distinct expression of key markers relating to thyroid hormone synthesis, including *TPO* and *DUOX1*. We also detected the expression of *ITLN1* and *MFGM*, which showed potential immune capacity and homeostasis maintenance of the epithelial tissue (Fig. 4b, Extended Data Fig. 6a). The expression of the key marker, *TPO*, was verified with ISH and the result showed a consistent expressional pattern with the previous report^14^ (Fig. 4c, Extended Data Fig. 6b). The ultracellular morphology of the cell clusters revealed that the region was composed of a single layer of non-ciliated cuboidal epithelial cells (Fig. 4d). To uncover the potential function of the region, expression profile-based functional pathway analyses were conducted based on the expression profile of the TLC. The expression score in functional pathways of thyroid hormone generation and metabolic process were projected on the spatial landscape showing intensive signals in zone 7 (Fig. 4e, Extended Data Fig. 6c).

To address the evolutionary homolog of the TLC to the thyroid gland, we conducted a cross-species comparison between the TLC cluster to the zebrafish developmental lineage/organs single-cell transcriptomic data. The result showed that the TLC had the highest similarity with the thyroid, liver, and pharyngeal endoderm cluster (Fig. 4f), verifying a strong expressional correlation between the TLC of the endostyle and pharyngeal endoderm (embryonic lineage) or thyroid (functional homolog) in zebrafish. Based on the result of gene markers detection, ISH verification, and cross-species comparison, we defined the TLC as a cell population with thyroid-equivalent features in ascidian endostyle, which verified the reliability of the cell atlas based on the previous report thyroid-equivalent region in the endostyle^19^.

To explore the potential developmental status and renewal capacity of the thyroid equivalent cell cluster, we further conducted the developmental trajectory assignment on cells in TLC. We first constructed a developing lineage with pharyngeal endoderm and the thyroid from the zebrafish, then obtained vital genes for thyroid gland developmental trajectory, which enriched in functional pathways including ECM-receptor interaction (Extended Data Fig. 6d-f). The expression level of vital thyroid developmental gene homologs in TLC were visualized and cells in TLC were assigned with pseudotime value based on the status of top expressed genes. Pseudotime frequency distribution showed cells in TLC are of diverse maturation status. Moreover, projection of pseudotime value on the cell atlas showed an enrichment of highly differentiated cells in dorsal region of the region (Fig. 4g). These results suggested a renewal trend or a differentiation trajectory of TLC, which is from ventral to dorsal.

## Discussion

The evolution and origination of complex organs have long been an intriguing topic in evo-devo since Charles Darwin’s emphasis on the significance of embryology in understanding evolution^52^. With the advent of multi-omics techniques such as single-cell and spatial transcriptomics, the comprehension of cell type composition and cell-to-cell interaction from the evo-devo perspective is rapidly advancing, providing new opportunities to investigate the evolution of organs in animals of significant evolutionary status^25^. By utilizing cutting-edge Stereo-seq and single-cell RNA-seq, we constructed the first spatially-resolved cell atlas in single-cell resolution for a distinct pharyngeal organ in basal chordates, the endostyle. This research innovatively provided the first profile of a marine animal spatial-resolved cell atlas. The assignment of transcriptomic profiles with spatial location has enabled the provision of locational information on transcriptional profile, which is vital in resolving organ composition with in complex structural organization and tissue arrangement patterns, compared to previous single-cell atlas datasets on marine animal samples^26, 53–55^.

Marine organisms exhibit exceptional variability across different species. The sample preparation method for spatial transcriptomics was optimized based on the previous study using fixed and embedded methods^56–58^ for ISH and single-molecule RNA-FISH. To ensure the replicability of the atlas, we combined three bio-replicons of single-cell RNA-seq and six Stereo-seq sections. All of these data show similar expression patterns, except for the statistical differences between two types of techniques (Extended Data Fig. 3b-3d). We interpretated the data from various bio-replicons and technical batches not only separately but also in combination, which ensured the reliability of our conclusions. To improve the alignment efficiency of transcriptomic data, we adjusted the genomic structural annotation and replenished alternative splicing information based on the previous publication^31^ (Supplementary Data 6), which largely improved the mapping results of transcriptional reads. The updated version has been released on EDomics, a multi-omics database for animal evo-devo^59^. Single-cell RNA-seq and Stereo-seq obtained 18,685 and 11,912 genes, respectively, compared to bulk RNA sequencing result (11,083 genes from bulk RNA sequencing^33^) (Extended Data Fig. 3g-h and Supplementary Data 5), which to some extend exhibit the difference between different techniques. Although the spatial transcriptome still has limited capture efficiency compared with single-cell RNA-seq, it shows irreplaceable advantages such as providing spatial information and having less bias in capturing rare cell types and signatures. Our results provide a detailed reference for *de novo* construction of cell atlas in marine species with limited prior research.

Besides zone 3 and zone 7, we conducted a detailed investigation of the cellular component and expressional features in DTR regions (Extended Data Fig. 7a-c). Zone 1 and 8 were composed of flagellate cell and ciliated cell clusters, both of which exhibited similar expression features related to ciliated cellular structures. Zone 2 and 4 shared similar morphology and expression, which were defined as PPIB- and PPIB+ secretory cells. Cell components in zone 6 were defined as the antimicrobial epithelium based on *ITLN2* expression. While information on zone 5 is still lacking due to its small tissue area as a connective tissue. Combined with the cellular components in HLR (Fig. 2a) and supported by previous studies on colony ascidian^20, 21^, the regional cell component of the endostyle was comprehensively uncovered. The cell populations defined in this atlas are not only consistent with previous understanding of the endostyle regions^15^, but also greatly enhance the comprehension on the pharyngeal transitional organ.

Based on the transcriptional cell atlas, we conducted a comprehensive comparison between the endostyle with organs/developmental lineages in zebrafish, revealing a high level of expression similarity of between the endostyle and multiple tissues in zebrafish. Interestingly, the endostyle displayed a particularly high similarity to myeloid lineage, which parallels previous discoveries of hematopoietic niche in the endostyle of colony ascidian^20^. This finding might suggest the existence of the hematopoietic niche in pharyngeal organ of vertebrate sister animal groups. In addition to the previously believed high expressional similarity to thyroid and pharyngeal endoderm, we also observed high expressional similarity to placode, integument and cochlea, indicating potential functional homolog between the ectoderm-derived tissue and the endostyle. These findings point to new research directions for understanding the evolution of pharyngeal organs.

In this study, we characterized the tissue continuity of the epithelial region extending from zone 9 and embracing an enclosed region that was previously overlooked as a blood sinus region. This region, which was defined as the HLR, was characterized by enriched blood and immune cell populations, potential developmental lineages, and highly developed cave-like ECM structures. The highly developed ECM may provide a favorable microenvironment for cell proliferation and lineage construction. These findings supported previous discoveries of hematopoietic niche^20^ and stem-cell harbor feature^21^ in the ‘sinus’ region, and further expand our understanding of developmental lineage of cell populations with stem capacity, such as IRF+ cells. The HLR is likely to be a primitive form for the blood and lymphoid system in vertebrates, including blood and immune organs like thymus and tonsils (Fig. 5b). However, the mixture of blood and immune system in HLR is strongly contrasting to the highly developed erythroid and myeloid cell lineage originating from hematopoietic stem cells in vertebrates.

The inner ear is believed a vertebrate innovation absent in non-vertebrate chordates and invertebrates^51^. Although, a variety of hair cell populations with potential mechanical forcing sensing function extensively exist in invertebrates^60, 61^. The direct evolutionary foundation for vertebrate inner ear still lacks a robust conclusion even some fragmentary evidences were reported in basal-chordates^62^. For example, the potential mechanoreceptors in amphioxus similar to neuromasts, are scattered over the body surface and unlikely to be homologous to vertebrates hair cells^63^. The potential hair cell populations in ascidians, including the primary sensory cells (ciliated-based cells)^64, 65^ and secondary sensory cells represented by coronal organs (form synapses with base nerve)^66^, have well-described morphology^67, 68^. However, they lack reasonable similarity to the body plan of vertebrate inner ear. In vertebrates, the inner ear is originated from otic vesicle initially located posterior to the dorsal portion of the second pharyngeal arch and involved fusion with the first and second arch originated middle ear. This process is reminiscent of pharynx-related components in the basal chrodates. Hence, evidence points to the existence of a functional hair cell population in cell population of the specialized pharyngeal organ endostyle. Strikingly, we noticed the high morphological similarity between zone 3 in endostyle to Kölliker’s organ, a critical epithelial structure in developing mammal auditory sensory organ^69^. We promote the preliminary conclusion that the hair cell cluster in the endostyle is likely the most ancient evolutionary cellular foundation for the vertebrate inner ear. While the relationship between the cupular sense organs^64^ and the hair cell population in endostyle remains further elucidation in the closest living relative of vertebrates, tunicates^70^.

The endostyle has long been believed developed from ventral pharyngeal endoderm^11, 13, 71^. Recent studies on both the oikopleura^72^ and *Ciona*^73^ showed the persistent development of supporting regions under the disturbance of Nkx2.1, which concluded that the supporting components of the endostyle are established parallel to regulation by other transcriptional factors. A series of markers were uncovered as hair cell architecture maintenance and neural transduction functions serving potential mechanical sensing in zone 3. Cross-species comparison reveals highly similarity between the cell population in zone 3 and acoustic-lateralis system in vertebrates, a typical ectoderm-origin tissue. The potential involvement of non-endoderm component in the endostyle development remains further investigation.

The origination of a key endocrine endoderm gland, the thyroid organ, can be traced back to cell clusters in the endostyle, which were believed to be capable of concentrating iodine and secreting thyroid hormone. In the ascidian endostyle, zone 7 was confirmed as the thyroid equivalent region from gene expression level^14^. In this study, we identified a cell clustered dominating zone 7 and defined it as thyroid like cell based on levels of evidence. Interestingly, pseudotime trajectory assignment with the developmental trajectory of thyroid gland in zebrafish showed the diverse developing status of TLC. Cells in various developing status co-exist in the region and statistical analysis exhibits an evolving trend of maturation tendency from the ventral to the dorsal region. This result showed that TLC may consistently replenish new mature cells or have the capacity of regeneration in response to tissue damage caused by damage. However, the divergence of TLC in ascidian is contrast with the very limited renewal capacity of thyroid gland in adult vertebrates^74^, which might provide a potential model for study in thyroid regeneration.

## Summary

In this research, targeting at uncovering the evolution of multi-germ layered complexed organs in vertebrate pharynx, we provide a high-quality dataset of a transitional pharyngeal organ that exclusively exists in basal chordates and proved its diverse cellular composition and potential evolutionary cellular prototypes of advanced pharynx organs in vertebrates. The HLR with immune/blood cell stemness and developmental trajectory is a potential homolog to advanced immune organs, including the tonsil and thymus. And cell populations in zone 3 are homolog candidates to auditory/lateralis system in vertebrates. We also verified the thyroid like cell cluster as a prototype of the thyroid gland in vertebrates and proved its self-renewal capacity. These discoveries strengthen the idea that endostyle is a key transitional organ in process of pharynx organ evolution with diverse functions homolog to advance vertebrates. The involvement of multiple cellular components in the pharynx of the basal chordate provides a cellular foundation of multiple advanced organs in vertebrates and illuminates the evolutionary footprint of pharyngeal architecture. Our research on pharyngeal organ evolution contribute to the understanding and investigation of new treatments for diseases in complex organs in humans.

## Methods

### Animals and sample collection

Adults *S. clava* were purchased from the Xunshan Company in Weihai, China and temporarily preserved in the aquarium system of laboratory at 18 °C with a consistent oxygen supply. The endostyle of healthy individual was gently dissected with disinfected instruments as previous description^33^.

### Single cell isolation and single-cell RNA-seq library construction

Fresh endostyle was transferred into clean vials and immediately dissociated using 0.2% trypsin in Ca^2+^-free ASW with 5 mM EGTA. The tissue was scissored and pipetted for 15-20 min on ice to completely dissociate the endostyle into individual cells. Digestion was inhibited using 0.2% BSA in Ca^2+^-free ASW. Cells were collected via centrifugation at 4 °C and 500 g for 2-5 min and resuspended in ice-cold Ca^2+^-free ASW containing 0.1% BSA. Single-cell gel beads in emulsion were generated using a Chromium Controller instrument (10× Genomics). Sequencing libraries were prepared using Chromium Single-Cell 3′ Reagent Kits (10× Genomics) according to the manufacturer’s instructions. After performing cleanup using a SPRIselect Reagent Kit, the libraries were constructed by performing the following steps: fragmentation, end-repair, A-tailing, SPRIselect cleanup, adaptor ligation, SPRIselect cleanup, sample index PCR, and SPRIselect size selection. Libraries were sequenced using Illumina NovaSeq system.

### Stereo-seq library preparation and sequencing

For Stereo-seq cryosection, endostyle sample from one single adult animal was isolated and immediately sink into methane pre-cooled under -20℃ for 5 min. Tissue under through rapid fixation were snap-frozen in Tissue-Tek OCT (4583, Sakura, Torrance, CA) with liquid nitrogen prechilled isopentane, then transferred to a -80℃ refrigerator for storage before cryosection. Before sectioning, the OCT-embedded tissue was placed into a -20℃ freezing microtome for equilibration 30 min. Cryosections were collected at 10 µm intervals along the transverse plane in a Leica CM1950 cryostat.

Stereo-seq experiments were performed as previous described by Chen et al.^29^. Stereo-seq chip was washed with NF-H2O supplemented with 0.05 U/ul RNase inhibitor (NEB, M0314L). Six sections from the anterior half endostyle were mounted onto a single Stereo-seq chip with 3-minute incubation on a Thermocycler Adaptor at 37℃. Then sections fixation was performed using methanol and then incubated for 40 min at -20℃. Sections on the chip were then stained with nucleic acid dye (Thermo Fisher, Q10212) for single-stranded DNA (ssDNA) imaging, and scanned using the Motic Custom PA53 FS6 microscope system to capture the full structure images (objective 20×).

Tissue sections were permeabilized in 0.1% pepsin (Sigma, P7000) with 0.01M HCL buffer and incubated at 37℃ for 6 minutes. The released and captured RNA on the Stereo-seq chip were reversely transcribed at 42℃ overnight using SuperScript II reverse transcription (RT) mix. Sections were washed twice with Wash Buffer and digested with Tissue Removal buffer (10 mM Tris-HCl, 25 mM EDTA, 100 mM NaCl, 0.5% SDS) at 37℃ for 30 minutes. The remaining RT products were collected and amplified by KAPA HiFi Hotstart Ready Mix (Roche, KK2602) with 0.8 µM cDNA-PCR primer. Sequencing libraries were prepared with PCR products following the steps of concentration quantification, DNA fragmentation, PCR amplification and purification. The purified PCR products were used for DNB generation and finally sequenced on MGI DNBSEQ-T1 sequencer (35 bp for Read1, 100 bp for Read2).

### Data analysis for single-Cell RNA-sequencing

For single-cell RNA-seq result from 10x Genomics Chromium, three samples were preprocessed respectively. BCL files were converted into FASTQ format using bcl2fastq v1.8.4 (Illumina). Cell Ranger v3.0.2 was used to generate gene-barcode matrix for each sample with mkref and count functions on default parameters. Reference genome were published in the previous research^31^. Seurat 4.0^75^ were used to construct analysis object based on expression matrix. Filtering threshold on the nCounts was from 0 to 5000, the nFeature was from 300 to 1200. Mitochondrial percent was calculated by dividing the mitochondrial gene counts the sum of count for a single cell.

Filtering threshold for mitochondrial expression percent were set to 30, 45, 35 for three samples respectively. Filtered data were processed with SCTransform method, variable features were 2000, min cells were 5 and method was qpoisson. Potential doublets in dataset were removed with DoubletFinder v3.0^76^, in which the parameter was 0.075 for assuming doublet formation rate and the pc was 15. SoupX^77^ were used to eliminate the ambient RNA in the buffer system, after which the data were processed with SCTransform again. Three filtered single-cell datasets were integrated with Seurat methods IntegrateData to remove batch effects. The integrated dataset was process by principal component analysis with default parameters. And the resolution was set to 0.3 for clustering. The marker genes for each cell cluster were identified using the FindAllMarkers function.

### Stereo-seq data analysis

Spatially-resolved RNA-seq data produced by Stereo-seq were pre-processed to generate gene expression matrices for subsequent analyses. The sequenced reads were mapped to their spatial location and corresponding gene structure using Stereo-seq Analysis Workflow (SAW, https://github.com/BGIResearch/SAW). Spatial gene expression matrices were generated using clean exon data only.

To tracing the cell to which each DNB data belong in the final spatial gene expression matrices, we performed object segmentation based on the nuclei in ssDNA staining image and then transferred the object outliner to the aligned gene expression spatial image. To do this, image registration was first performed to align the imaging ssDNA to the coordinates of gene expression matrices. The spatial gene expression matrices were converted into 2D spatial image by color-coding the UMI counts of each DNB. While the corresponding ssDNA image was processed following steps of background noise filtering, local contrast enhancement, and then transformed into their real physical size by replacing the pixel unit with the distance of DNBs (715 nm). Then the pre-processed image datasets were aligned with the TrakEM2 algorithm by Fiji, in which the tissue morphology was manually detected and used for feature matching. After the affine transformation, the ssDNA images were used for object segmentation with the CellProfiler software^78^. Briefly, nuclei location was detected with the global Otsu threshold strategy without manual annotation, and the Mutex Watershed algorithm was used to automatically determine whether the pixel belongs to a specific nucleus. Then the RNA data of each DNB location was assigned into individual cell object based on whether they fell within the segmented cell outliners. As for the regions in which dense nuclei located and failed to detect the boundary, we manually masked these regions and retained bin15 (around 10 µm in width and height) partition instead of cell segmentation. With the aligned coordinates of cells and gene expression, the gene expression matrices were further aggregated into putative cells ready for downstream analysis.

The aggregated cell level spatial RNA data were normalized by the SCTransform function in Seurat 4.0 and were further integrated with object from different sections using IntegrateData function. PCA was performed with default parameters and the top 30 principal components were extracted for unsupervised clustering with resolution parameter of 0.8. Spatial information was not considered for the clustering algorithm and we projected the resultant Seurat clusters into their spatial coordinate based on the cell segmented location.

### Data integration of single-cell RNA-sequencing and Stereo-seq

Three biological replicons from single-cell RNA-seq and six Stereo-seq section data were integrated with Seurat IntegrateData function with SCT normalize method. The integrated data were processed with principal component analysis and clustered with resolution 0.8. Differential expressed markers were exported with the FindAllMarkers method, based on which the cell type was annotated manually.

To compare the similarity and difference of scRNA-seq data and Spatial data, we project annotations from each of them into the same integrated UMAP dimension and manually verified the consistence of these two datasets.

### Identification of spatially correlated modules

Gene modules were categorized with autocorrelation algorithm, hotspot, based on spatial location^34^. The spatial expression matrix was normalized by the total UMI number of each putative cell after filtering genes that expressed in less than 3 cells. K-nearest neighbor (knn) graph of genes was further created with parameter of 300 neighbors. Genes with significant spatial autocorrelation (FDR < 0.05) were kept for computing local correlations. The gene modules were identified using the create_modules function with minimal gene threshold of 20 and fdr threshold of 0.05.

### Selection of zebrafish single-cell transcriptomic dataset

To investigate how similar the endostyle cell types to vertebrate tissues, we collected scRNA-seq data of zebrafish from Farnsworth et al.^36^, Qian et al.^37^ and Gillotay et al.^38^ as reference datasets. Before performing comparison of endostyle and zebrafish tissues, we pre-process the zebrafish datasets to unify the annotations and eliminate possible batches. These three zebrafish development and tissue specific single cell expression objects were separately normalized and then integrated using Seurat 4.0. PCA analysis was performed on the integrated assay and the top 30 principal components were selected for clustering. The integrated assay was then re-clustered with resolution of 0.8 to generate uniform clustering labels, after which cells with similar stat or identify should obtain the same label even they come from different data source. Annotations were manually updated based on the original cell types and our newly integrated clustering.

### Cross-species comparison between *S. clava* and zebrafish

Orthologs were identified using Blast for cross-species comparison. Coding sequences from both *S. clava* and zebrafish were aligned to each other using BLASTN. Matches with E-value less than 10^-6^ and identity value greater than 30% were kept as confident hits and further used in SAMap^79^ comparison. Only gene pairs with reciprocal match were identified as one-to-one orthologs for calculating correlation of the endostyle dataset and zebrafish dataset.

The correlation calculation was performed using Spearman algorithm based on the average expression of 387 homologous TF genes (Supplementary Data 4) in *S. clava* cell types and zebrafish tissues. Then we averaged the values of different cell types in *S. clava* as a tissue level correlation to that of each zebrafish tissue. To infer contributions of each kind of cell in endostyle to most similar tissues in zebrafish, we selected the top 50% zebrafish tissues and further estimated correlations between each endostyle cell type and zebrafish tissue. In this process, group of cells in developing phase as well as developed tissues were considered separately.

To compare specific endostyle cell type and zebrafish tissues, we subset the endostyle thyroid-like cells and zebrafish pharyngeal endoderm related groups, as well as endostyle hair cells and zebrafish hair cells related groups to apply SAMap analysis.

### Cell trajectory analysis of HLR cells

Monocle 3^80^ were utilized in constructing the pseudotime trajectory for the HLR cells which we speculated hierarchy lineage exists based on the expressed signatures. Only cell types that primarily located in the HLR region were kept and used for the construction of trajectory, and cells located in the DTR region were further filtered. The subset data from each section was process with Seurat 4.0 and integrated into a single object. The Seuratwrapper were used in connecting the Seurat results for Monocle 3 further analysis. Methods includes learn_graph and order_cells were used to construct the trajectory. The root of trajectory was assigned as cell type which nearest neighbored to ‘stem cell’ from single-cell RNA-seq dataset in our UMAP space.

Dynamo^81^ analysis was also performed follow the instructions at https://dynamo-release.readthedocs.io/. The non-spliced and spliced raw count matrices were calculated by Velocity package based on the annotation file and bam file generated by SAW, and were further processed by recipe_monocle function in Dynamo. Top 3000 highly expression genes were selected to perform dimension reduction via UMAP with default parameters. RNA velocity vectors were estimated on the normalized matrix and further were projected into the spatial visualization.

### Cell-cell interaction analysis

The ligand-receptor pairs from CellChatDB^82^ were used to evaluate cell-cell interaction between each pair of cell types. The gene homologous information of each ligand or receptor gene to that of *S. clava* were conducted with all protein sequences of two species with SonicParanoid^83^ software. All ortholog genes, including one-to-one and one-to-many, were used as possible matches to further infer ligand-receptor relationship in *S. clava*. Interaction score was calculated using cell2cell^84^ package following the instructions at https://earmingol.github.io/cell2cell. Target cell types, here hair cells and IRF positive cells, were selected by manually determining whether they are spatially neighbored.

### Cross-species trajectory projection

To investigate the evolutionary location and developmental stages of endostyle thyroid-like cells (TLC), we evaluated the similarity of TLC to different trajectory points of thyroid development, and project the TLC into a pseudo-evolutionary trajectory. We first constructed the pseudotime trajectory of pharyngeal endoderm and thyroid with Monocle 2^85^ to tracing the developmental footprint of zebrafish thyroid. Highly expressed genes (log_avg2FC >= 0.25) in pharyngeal endoderm and thyroid were estimated using differentialGeneTest function to find genes that change along pseudotime. By identifying the pseudotime point with maximal expression, these trajectory-related genes were then classified into different clusters and were treated as representation of developmental stages. Finally, we ordered the endostyle TLC cells based on their highest expressed trajectory-related genes and inferred the order as the pseudo-trajectory of endostyle thyroid like cells.

### ISH on the endostyle and peripheral blood

The fresh endostyle sample was fixated by 4 % paraformaldehyde (PFA) in artificial water at 4 °C for overnight and finish sucrose sinkage with 30 % sucrose in PBS solution. The dried tissue was embedded with O.C.T. compound (Sakura 4583) and snap-freeze with liquid nitrogen. Cryosections for ISH were prepared with Leica CM3050 and sections were sticked on to adhesive slides. The peripheral blood of *S. clava* were obtained from the non-endostyle pharynx blanket tissue and fixed with 4 % PFA. Blood cell drops washed by PBS were dried on the adhesive slides. Digoxigenin (DIG)-RNA probes were designed (Supplementary Table 7) and synthesized by in vitro transcription with SP6 RNA polymerase and T7 RNA polymerase (Thermo Fishers, EP0131, EP0111). The ISH procedure was conducted in the humified black box preventing the slide from drying. Regions with tissue on slides were circled with hydrophobic paint. The tissue was digested with 10 μg/mL Protenase K for 30 min at 37 °C and washed with PBST (0.1 % Tween 20 in PBS). Afterwards the tissue was refixed with 4% PFA in PBS solution for 30 min at room temperature and washed with PBST. The tissue was then incubated with pre-hyb buffer (50% formamide, 5x saline sodium citrate buffer (SSC), 0.1% Tween 20) at room temperature for 10 min. And the tissue was incubated with hybrid buffer (50% formamide, 5x SSC, 0.1% Tween 20, 5x Denhardt’s solution, 100 μg/mL yeast tRNA, and 50 μg/mL heparin) at hybrid temperature for 4 hrs. Then add new hybrid buffer with probes at concentration of 0.5 ng/μg at hybrid temperature for 20 hrs. Tissue were washed with (50% formamide, 5x SSC, 0.1% Tween 20) for four time and PBST for four times. Next, tissue was incubated in 1% blocking buffer (Roche) with Dig-antibody at 4 °C overnight. Finally, tissue was incubated with NBT-BCIP for chromogenic detection.

### Cytoskeleton and nuclear staining of transverse sections

Cryosections were prepared as the protocol mentioned above. The slide was incubated with 0.1% PBST (Triton X 100) for 20 min at room temperature. After which the slide was washed with PBS solution for three times. Dilute the Phalloidin 488 (Alexa, A12379) with PBS with proportion of 1:100 and stained the slide avoid light at room temperature for 30 min. Seal the slide with VECTASHIELD antifade mounting medium with DAPI (H-1200-10) and resin. Prepared slides were visualized with confocal microscope ZEISS LSM 900.

### TEM sample preparation

Fresh endostyle were collected and immersed into the fixation buffer (2.5 % glutaraldehyde in artificial seawater) and preserved under 4 °C. The semithin section and ultrathin section were prepared by Biomedical center of Qingdao medical college, Qingdao University.

### Statistics analysis

All the statistical analyses were performed using R (version 3.6). Student’s *t* test and Spearman’s rank correlation analysis were utilized in this study. The *P* value < 0.04 was considered as significant difference exists. * Represents 0.01 < *P* < 0.05. ** Represents 0.001 < *P* < 0.01. *** Represents *P* < 0.001.

## Data availability

All data generated or analyzed during this study are included in the manuscript and supporting files. The Stereo-seq and single-cell RNA-sequencing datasets generated in this study have been deposited at China National GeneBank DataBase (CNGBdb, https://db.cngb.org/) under the accession number CNP0004228.

## Author contributions

B.D., G.F., A.J., and J.W. conceived the idea; B.D., and G.F. supervised the work; W.Z. and J.W. generated the Single-cell RNA-seq library; X.S., X.L., A.J. and N.Z generated the Stereo-seq library; K.H. performed data preprocessing and quality evaluation; K.H., A.J., R.W. analyzed the data; J.Q., P.L., J.Z., Y.G., and Y.Z. provided technical and experimental support; Q.L, H. Y., S. W., K. C., X. X., and H. Y gave the relevant advice; A.J., J.W., K.H., and B.D. wrote and revised the manuscript.

## Acknowledgements

This work was funded by the Science & Technology Innovation Project of Laoshan Laboratory (No. LSKJ202203002), the National Key Research and Development Program of China (2022YFC2601302), and the Taishan Scholar Program of Shandong Province, China (B.D.).

---

**Extended Data Fig. 1:**
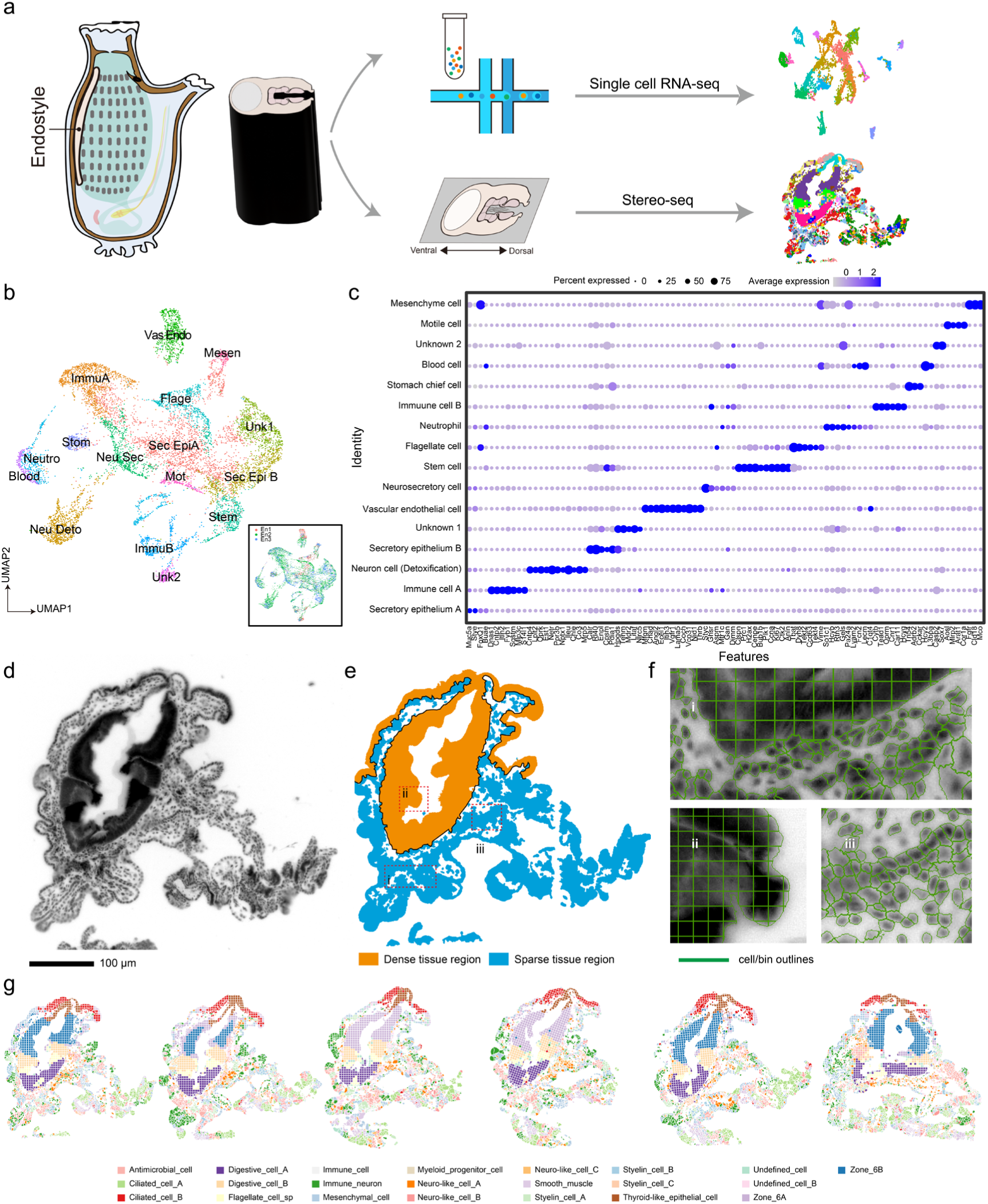
Workflow and strategy for the endostyle cell atlas construction. (a) Basic workflow for data acquisition. The endostyle of ascidian were manually dissociated and processed single-cell RNA-seq and stereo-seq. Libraries constructed in two techniques were sequenced and downstream analysis were conducted. (b) UMAP for cell clusters with cell type annotation by single-cell RNA-seq. Three biological replicons are implicated in the corner figure. (c) Marker genes in annotating cell clusters in the dataset of single-cell RNA-seq. (d) The optical image captured during library construction. The tissue was stained with ssDNA dye. (e) Two strategies in cell segregation. Dense tissue region was selected manually based on the optical image, which is segregated with the bin 15 division method. Surrounding region were segregated with Cellprofiler. (f) Segregated cell boundary for squared bin and segmented cell. (g) Cell atlas of six sections in Stereo-seq, colored with cell annotation on Stereo-seq dataset.

**Extended Data Fig. 2:**
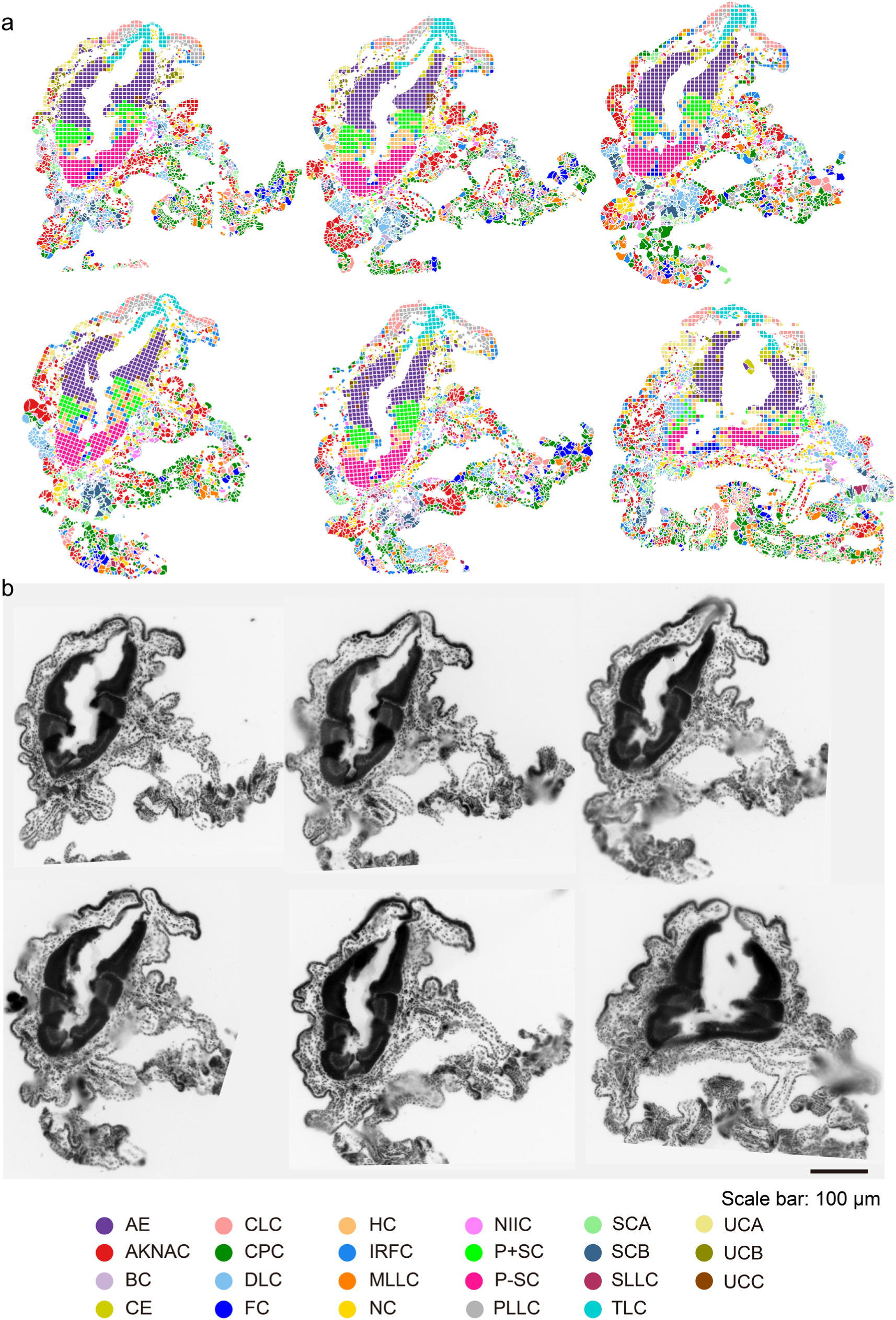
Cell population distribution and morphology correspondence in six sections of the endostyle cell atlas. (a) Cell population distribution in six sections. (b) ssDNA staining result for six sections. Bar: 100 μm.

**Extended Data Fig. 3:**
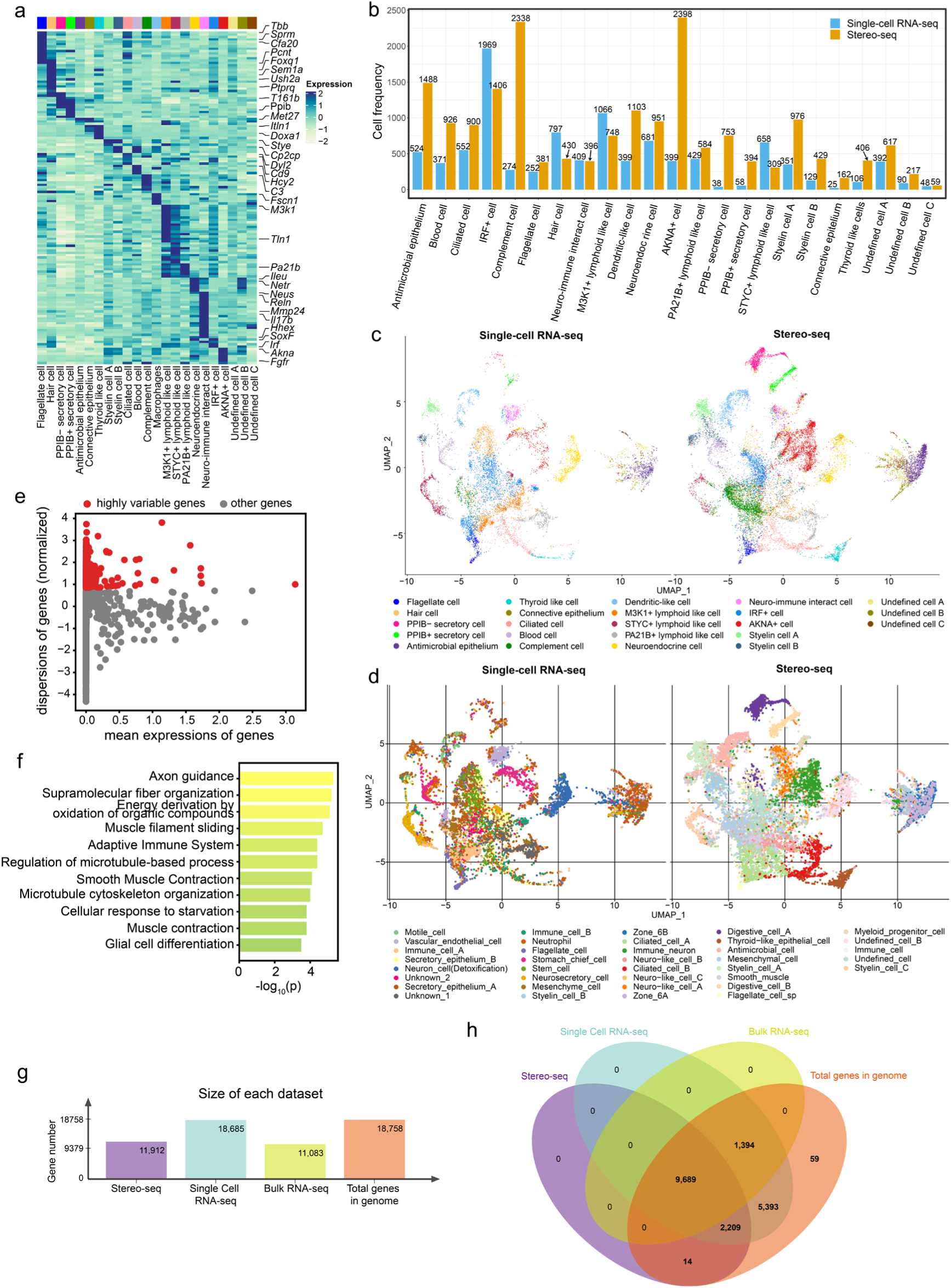
Analysis on the integrated dataset and two technical batch comparison. (a) Heatmap showing key marker genes in 23 cell clusters in the integrated dataset. Color bars on the head of each column is UMAP color code. (b) Bar plot showing cell frequencies of two technical batches in cell clusters. Blue bars are data from single-cell RNA-seq and orange bars are data from Stereo-seq. (c) UMAP separately showing the integrated dataset from two technical batches. Cell units were colored with cell type annotations on the integrated dataset. (d) UMAP separately showing the integrated dataset from two technical batches. Cell units were colored with cell type annotations on two cell batches respectively. (e) Dot plot showing the degree of gene expressional variation. Highly variable genes were colored with red, which are selected based on dispersion level of genes. (f) Bar plot showing the GO terms obtained from GO enrichment analysis on highly variable genes. (g) Number of detected genes in technical batches contrasting with the total gene number in genome. (h) Venn diagram of detected genes in technical batches.

**Extended Data Fig. 4:**
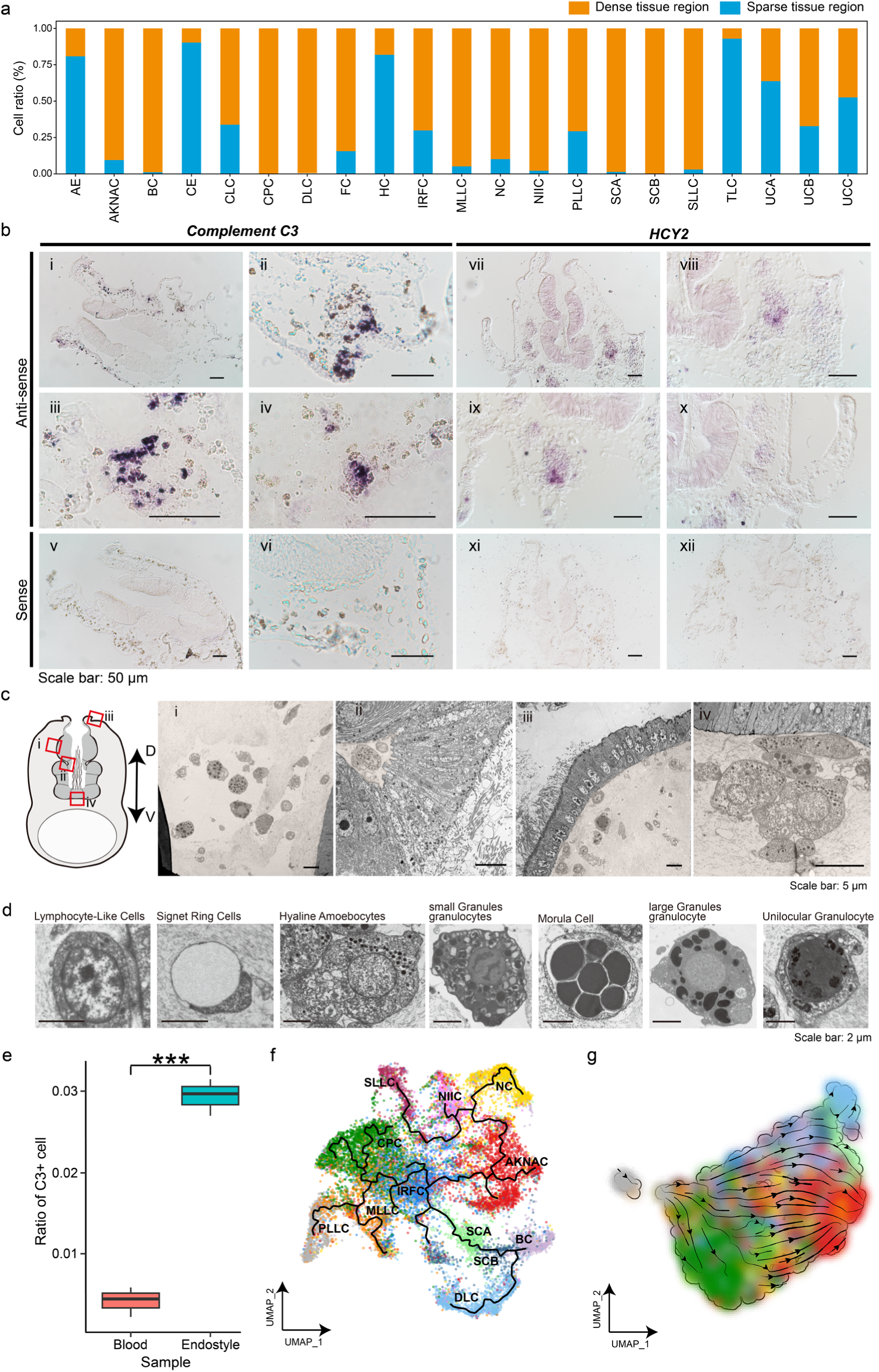
The HLR cell population and trajectory information. (a) Cell ratio distribution of each cell type in dense tissue region and sparse tissue region. (**b**)ISH of *C3* and *HCY2* on the endostyle cryosection. (i-iv) and (v-vi) showing antisense and sense results for *C3*. (vii-x) and (xi-xii) showing antisense and sense results for *HCY2*. (c) Left, schematic for the location of the HLR detailed structure. Right, Red rectangle regions for TEM visualization in (i-iv). HLR in TEM pictures were masked with light pale pink. (d) Morphology of distinguished blood cell type in the HLR. (e) Statistical analysis for the ratio of C3+ cells in the endostyle HLR and peripheral blood respectively. *** *P* < 0.001. (f) Pseudotime trajectory analysis depicting trajectories among cell clusters in the HLR based on the UMAP. Cells are colored and labeled with cell type annotations. The trajectory is plotted with curved lines. (g) RNA velocity streamline plots showing the predicted trajectory of cell clusters transition in the HLR of the endostyle.

**Extended Data Fig. 5:**
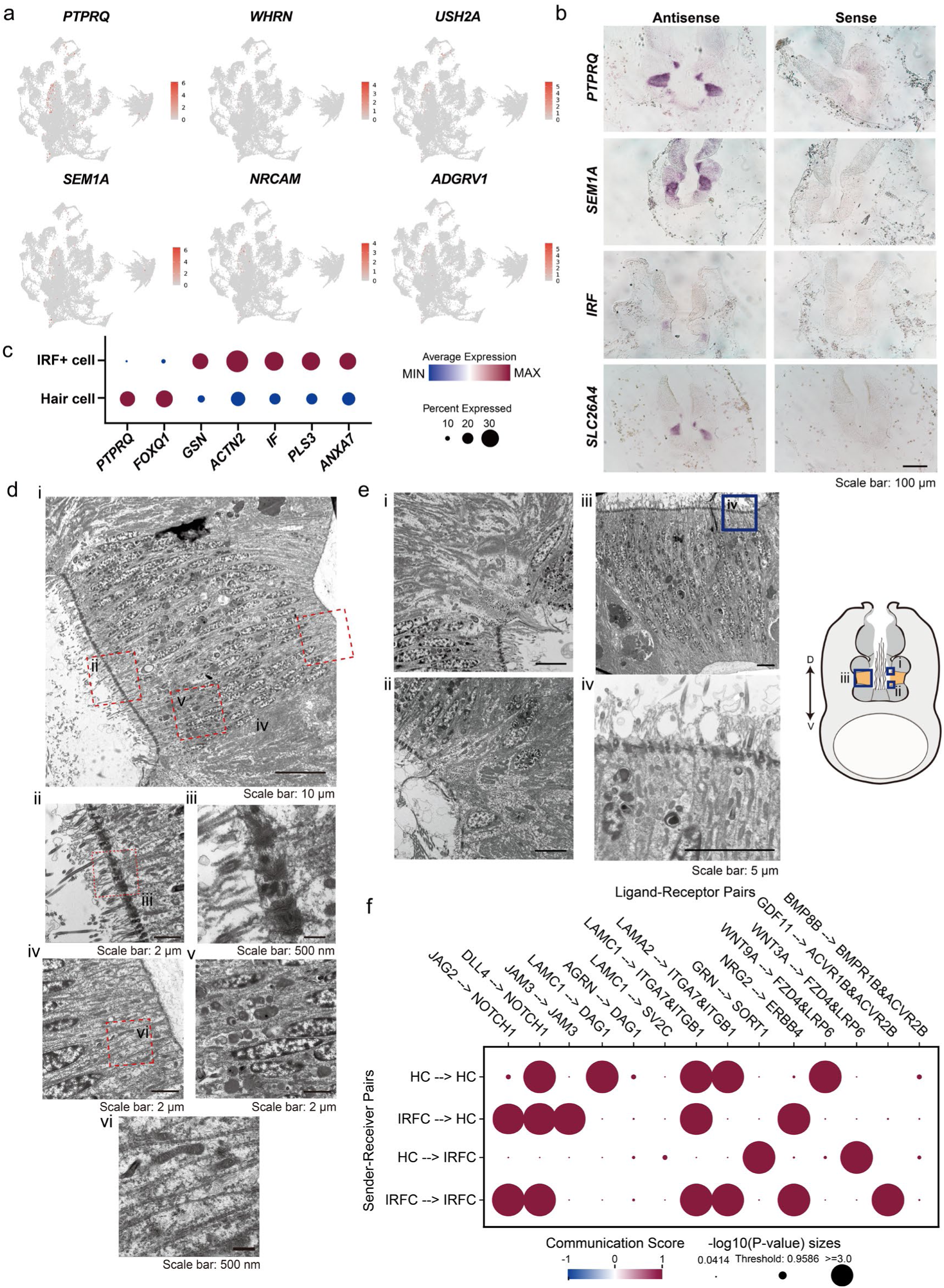
Expression pattern and morphology of hair cell candidate in zone 3. **(a)**UMAP showing the expression of key markers in hair cell candidate in zone 3 of the endostyle. The expression levels of marker genes were projected on the UMAP for the integrated dataset. (b) ISH showing the expression pattern of the *PTPRQ*, *SEM1A*, *IRF* and *SLC26A4* genes. (c) Bubble plot showing the expression of distinct expressed gene for the hair cell and IRF+ cell clusters. (d) Morphology observation with TEM showing the detailed structure in zone 3. (i) Whole scene of zone 3 in the endostyle. (ii) Detailed structure in red rectangle region of the apical surface of zone 3 in (i). (iii) Detailed structure in red rectangle region in (ii). (iv) Detailed structure of red rectangle region in (i) basal region. (v) Detailed structure of red rectangle region in (i) within tissue. (vi) Detailed structure of red rectangle region in (iv). (e) Morphology observation with TEM showing the boundary of zone 3 with zone 2 and zone 4. (i) Detailed structure for the boundary region of zone 4 and zone 3. (ii) Detailed structure for the boundary region of zone 2 and zone 3. (iii) Apical region of zone 3. (iv) Detailed structure of blue rectangle region of (iii). D: dorsal, V: ventral. (f) Bubble plot showing communication scores of ligand-receptor pairs between sender and receiver cell populations.

**Extended Data Fig. 6:**
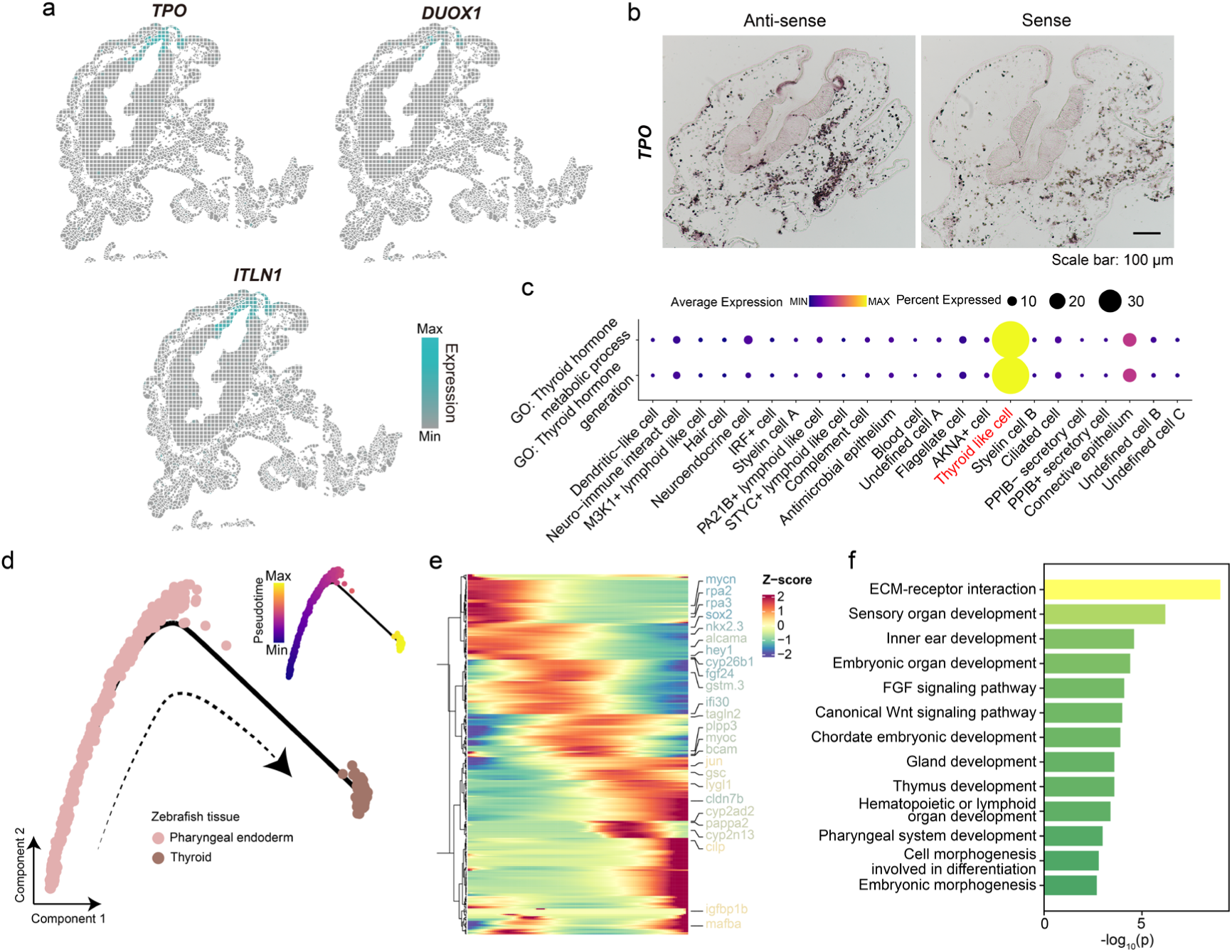
The thyroid like cell cluster expression features and functional analysis. (a) Heatmap showing expression patterns of key markers on spatial landscape. (b) ISH showing the expression pattern of *TPO* on the endostyle transverse section. (c) Bubble plot showing the expression of genes on function pathways. (d) The cell trajectory of pharyngeal endoderm and thyroid from single-cell RNA-seq data of zebrafish. (e) Gene expression cascades during the development of the thyroid gland from the pharyngeal endoderm. The gene names were colored according to their gene trajectory status, which suggesting their expression variation trend in pseudotime trajectory. (f) Barplot showing the GO analysis results of the functional terms in the gene set of (e).

**Extended Data Figure 7:**
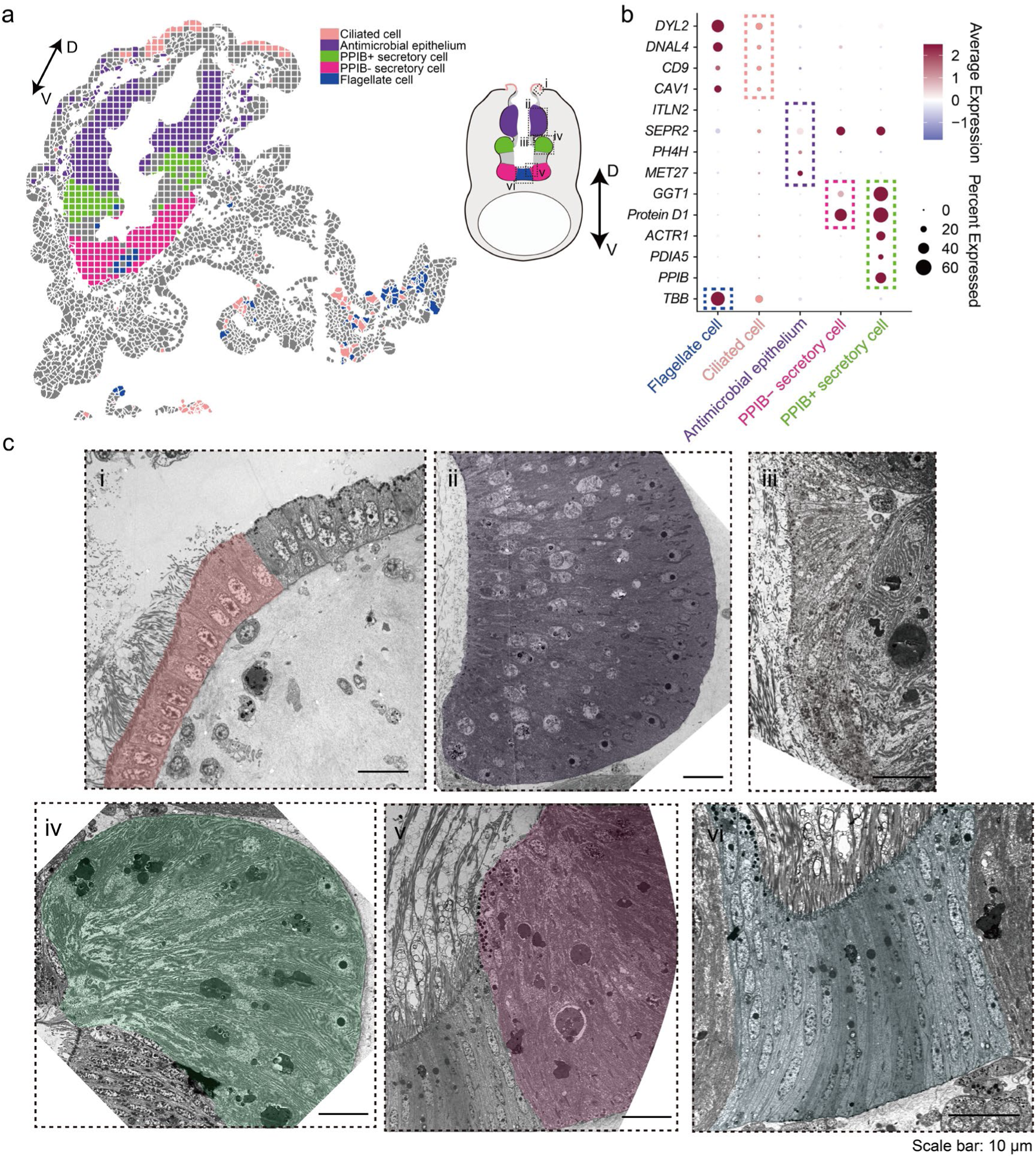
The expression and morphology of DERs in the endostyle. (a) Spatial visualization of ciliated cell. Antimicrobial epithelium, PPIB+ secretory cell, PPIB-secretory cell and flagellate cell clusters, corresponding to main cellular composition of zone 8, 6, 4, 2, 1. D: dorsal, V: ventral. (b) Bubble plot showing expression pattern of marker genes in indicated cell clusters. Expression bubbles of marker for specific cluster were labeled with colored rectangular. (c) Ultracellular structures of indicated cell clusters in DERs. The tissue was labeled with corresponding color. (i) Morphology of simple epithelium in zone 8. Cells in zone 8 are highly ciliated, while the adjacent region, zone 9 are epithelium without cilia. (ii) Morphology of fan-shape epithelium in zone 6. Vacuoles can be seen in the apical side. (iii) Morphology of epithelium in zone 5, which is a connective region between zone 4 and zone 6. (iv) Morphology of fan-shape epithelium in zone 4. Cilia and particles can be seen in apical surface. Highly developed ER can be seen. (v) Morphology of fan-shape epithelium in zone 2. The morphology is similar to that of zone 4. (vi) Morphology of column epithelium with long cilia in zone 1.

## References

1 Gee, H. Across the Bridge: Understanding the Origin of the Vertebrates. (The University of Chicago Press, 2018).

2 Gillis, J. A., Fritzenwanker, J. H. & Lowe, C. J. A stem-deuterostome origin of the vertebrate pharyngeal transcriptional network. Proceedings of the Royal Society B: Biological Sciences 279, 237–246, doi:10.1098/rspb.2011.0599 (2012).

3 Lowe, C. J., Clarke, D. N., Medeiros, D. M., Rokhsar, D. S. & Gerhart, J. The deuterostome context of chordate origins. Nature 520, 456–465, doi:10.1038/nature14434 (2015).

4 Clausen, S. & Smith, A. B. Palaeoanatomy and biological affinities of a Cambrian deuterostome (Stylophora). Nature 438, 351–354, doi:10.1038/nature04109 (2005).

5 Tian, Q., Zhao, F., Zeng, H., Zhu, M. & Jiang, B. Ultrastructure reveals ancestral vertebrate pharyngeal skeleton in yunnanozoans. Science 377, 218–222, doi:10.1126/science.abm2708 (2022).

6 Pardos, F. Fine Structure and Function of Pharynx Cilia in Glossobalanus minutus Kowalewsky (Enteropneusta). Acta Zoologica 69, 1–12, doi:10.1111/j.1463-6395.1988.tb00895.x (1988).

7 Veitch, E., Begbie, J., Schilling, T. F., Smith, M. M. & Graham, A. Pharyngeal arch patterning in the absence of neural crest. Current Biology 9, 1481–1484, doi:10.1016/S0960-9822(00)80118-9 (1999).

8 Graham, A. & Richardson, J. Developmental and evolutionary origins of the pharyngeal apparatus. EvoDevo 3, 24, doi:10.1186/2041-9139-3-24 (2012).

9 Piotrowski, T. & Nusslein-Volhard, C. The endoderm plays an important role in patterning the segmented pharyngeal region in zebrafish (*Danio rerio*). Developmental Biology 225, 339–356, doi:10.1006/dbio.2000.9842 (2000).

10 Anthwal, N. & Thompson, H. The development of the mammalian outer and middle ear. Journal of Anatomy 228, 217–232, doi:10.1111/joa.12344 (2016).

11 Olsson, R. Endostyles and endostylar secretions: A comparative histochemical study. Acta Zoologica 44, 299–328, doi:10.1111/j.1463-6395.1963.tb00411.x (1963).

12 Petersen, J. K. Ascidian suspension feeding. *Journal of Experimental Marine Biology and Ecology* **342**, 127–137, doi:10.1016/j.jembe.2006.10.023 (2007).

13 Hirano, T. & Nishida, H. Developmental fates of larval tissues after metamorphosis in the ascidian, Halocynthia roretzi. II. Origin of endodermal tissues of the juvenile. Development Genes and Evolution 210, 55–63, doi:10.1007/s004270050011 (2000).

14 Ogasawara, M., Di Lauro, R. & Satoh, N. Ascidian homologs of mammalian thyroid peroxidase genes are expressed in the thyroid-equivalent region of the endostyle. Journal of Experimental Zoology 285, 158–169, doi:10.1002/(sici)1097-010x(19990815)285:2<158::aid-jez8>3.0.co;2-0 (1999).

15 Hiruta, J., Mazet, F., Yasui, K., Zhang, P. & Ogasawara, M. Comparative expression analysis of transcription factor genes in the endostyle of invertebrate chordates. Developmental Dynamics 233, 1031–1037, doi:10.1002/dvdy.20401 (2005).

16 Sasaki, A., Miyamoto, Y., Satou, Y., Satoh, N. & Ogasawara, M. Novel endostyle-specific genes in the ascidian *Ciona intestinalis*. Zoological Science 20, 1025–1030, doi:10.2108/zsj.20.1025 (2003).

17 Barrington, E. J. & Thorpe, A. The identification of monoiodotyrosine, diiodotyrosine and thyroxine in extracts of the endostyle of the ascidian, Ciona Intestinalis Linnaeus. Proceedings of the Royal Society B: Biological Sciences 163, 136–149, doi:10.1098/rspb.1965.0063 (1965).

18 Barrington, E. J. W. & Thorpe, A. An autoradiographic study of the binding of iodine125 in the endostyle and pharynx of the ascidian, *Ciona intestinalis L*. General and Comparative Endocrinology 5, 373–385, doi:10.1016/0016-6480(65)90062-6 (1965).

19 Nilsson, M. & Fagman, H. Development of the thyroid gland. Development 144, 2123–2140, doi:10.1242/dev.145615 (2017).

20 Rosental, B. et al. Complex mammalian-like haematopoietic system found in a colonial chordate. Nature 564, 425–429, doi:10.1038/s41586-018-0783-x (2018).

21 Voskoboynik, A. et al. Identification of the endostyle as a stem cell niche in a colonial chordate. Cell Stem Cell 3, 456–464, doi:10.1016/j.stem.2008.07.023 (2008).

22 Osugi, T., Sasakura, Y. & Satake, H. The ventral peptidergic system of the adult ascidian *Ciona robusta* (*Ciona intestinalis* Type A) insights from a transgenic animal model. Scientific Reports 10, 1892, doi:10.1038/s41598-020-58884-w (2020).

23 Parrinello, D., Sanfratello, M. A., Vizzini, A., Parrinello, N. & Cammarata, M. Ciona intestinalis galectin (CiLgals-a and CiLgals-b) genes are differentially expressed in endostyle zones and challenged by LPS. Fish & Shellfish Immunology 42, 171–176, doi:10.1016/j.fsi.2014.10.026 (2015).

24 Vizzini, A. et al. Upregulated transcription of phenoloxidase genes in the pharynx and endostyle of *Ciona intestinalis* in response to LPS. Journal of Invertebrate Pathology 126, 6–11, doi:10.1016/j.jip.2015.01.009 (2015).

25 Arendt, D. et al. The origin and evolution of cell types. Nature Reviews Genetics 17, 744–757, doi:10.1038/nrg.2016.127 (2016).

26 Cao, C. et al. Comprehensive single-cell transcriptome lineages of a proto-vertebrate. Nature 571, 349–354, doi:10.1038/s41586-019-1385-y (2019).

27 Khrameeva, E. et al. Single-cell-resolution transcriptome map of human, chimpanzee, bonobo, and macaque brains. Genome Res 30, 776–789, doi:10.1101/gr.256958.119 (2020).

28 Farnsworth, D. R., Saunders, L. M. & Miller, A. C. A single-cell transcriptome atlas for zebrafish development. Developmental Biology 459, 100–108, doi:10.1016/j.ydbio.2019.11.008 (2020).

29 Chen, A. et al. Spatiotemporal transcriptomic atlas of mouse organogenesis using DNA nanoball-patterned arrays. Cell 185, 1777–1792 e1721, doi:10.1016/j.cell.2022.04.003 (2022).

30 Macosko, E. Z. et al. Highly Parallel Genome-wide Expression Profiling of Individual Cells Using Nanoliter Droplets. Cell 161, 1202–1214, doi:10.1016/j.cell.2015.05.002 (2015).

31 Wei, J. et al. Genomic basis of environmental adaptation in the leathery sea squirt (*Styela clava*). Molecular Ecology Resources 20, 1414–1431, doi:10.1111/1755-0998.13209 (2020).

32 Zhang, J., Wei, J., Yu, H. & Dong, B. Genome-Wide Identification, Comparison, and Expression Analysis of Transcription Factors in Ascidian *Styela clava*. International Journal of Molecular Sciences 22, doi:10.3390/ijms22094317 (2021).

33 Jiang, A., Zhang, W., Wei, J., Liu, P. & Dong, B. Transcriptional Analysis of the Endostyle Reveals Pharyngeal Organ Functions in Ascidian. Biology 12, doi:10.3390/biology12020245 (2023).

34 DeTomaso, D. & Yosef, N. Hotspot identifies informative gene modules across modalities of single-cell genomics. Cell Systems 12, 446–456.e449, doi:10.1016/j.cels.2021.04.005 (2021).

35 Holley, M. C. Cell shape, spatial patterns of cilia, and mucus-net construction in the ascidian endostyle. Tissue & cell 18, 667–684, doi:10.1016/0040-8166(86)90069-8 (1986).

36 Tambalo, M., Mitter, R. & Wilkinson, D. G. A single cell transcriptome atlas of the developing zebrafish hindbrain. Development 147, doi:10.1242/dev.184143 (2020).

37 Qian, F. et al. Single-cell RNA-sequencing of zebrafish hair cells reveals novel genes potentially involved in hearing loss. Cellular and Molecular Life Sciences 79, 385, doi:10.1007/s00018-022-04410-2 (2022).

38 Gillotay, P. et al. Single-cell transcriptome analysis reveals thyrocyte diversity in the zebrafish thyroid gland. EMBO reports 21, e50612, doi:10.15252/embr.202050612 (2020).

39 Graham, A. The development and evolution of the pharyngeal arches. J Anat 199, 133–141, doi:10.1046/j.1469-7580.2001.19910133.x (2001).

40 Longo, V. et al. The conservation and diversity of ascidian cells and molecules involved in the inflammatory reaction: The *Ciona robusta* model. Fish & Shellfish Immunology 119, 384–396, doi:10.1016/j.fsi.2021.10.022 (2021).

41 Zhang, W. et al. Protein Tyrosine Phosphatase Receptor-type Q: Structure, Activity, and Implications in Human Disease. Protein & Peptide Letters 29, 567–573, doi:10.2174/0929866529666220511141826 (2022).

42 Castiglione, A. & Möller, C. Usher Syndrome. Audiology Research 12, 42–65 (2022).

43 Michalski, N. & Petit, C. Genes Involved in the Development and Physiology of Both the Peripheral and Central Auditory Systems. Annu Rev Neurosci 42, 67–86, doi:10.1146/annurev-neuro-070918-050428 (2019).

44 Mathur, P. D. & Yang, J. Usher syndrome and non-syndromic deafness: Functions of different whirlin isoforms in the cochlea, vestibular organs, and retina. Hearing research 375, 14–24, doi:10.1016/j.heares.2019.02.007 (2019).

45 Zhou, P. et al. ADGRV1 Variants in Febrile Seizures/Epilepsy With Antecedent Febrile Seizures and Their Associations With Audio-Visual Abnormalities. Frontiers in Molecular Neuroscience 15, doi:10.3389/fnmol.2022.864074 (2022).

46 Brand, Y. et al. Neural cell adhesion molecule NrCAM is expressed in the mammalian inner ear and modulates spiral ganglion neurite outgrowth in an in vitro alternate choice assay. J Mol Neurosci 55, 836–844, doi:10.1007/s12031-014-0436-y (2015).

47 Cafferty, P., Yu, L., Long, H. & Rao, Y. Semaphorin-1a functions as a guidance receptor in the Drosophila visual system. Journal of Neuroscience 26, 3999–4003, doi:10.1523/JNEUROSCI.3845-05.2006 (2006).

48 Wémeau, J. L. & Kopp, P. Pendred syndrome. Best Pract Res Clin Endocrinol Metab 31, 213–224, doi:10.1016/j.beem.2017.04.011 (2017).

49 Vullhorst, D. et al. A negative feedback loop controls NMDA receptor function in cortical interneurons via neuregulin 2/ErbB4 signalling. Nat Commun 6, 7222, doi:10.1038/ncomms8222 (2015).

50 Hu, F. et al. Sortilin-mediated endocytosis determines levels of the frontotemporal dementia protein, progranulin. Neuron 68, 654–667, doi:10.1016/j.neuron.2010.09.034 (2010).

51 Higuchi, S. et al. Inner ear development in cyclostomes and evolution of the vertebrate semicircular canals. Nature 565, 347–350, doi:10.1038/s41586-018-0782-y (2019).

52 Hall, B. K. Evolutionary Developmental Biology (Evo-Devo): Past, Present, and Future. Evolution: Education and Outreach 5, 184–193, doi:10.1007/s12052-012-0418-x (2012).

53 Ma, P. et al. Joint profiling of gene expression and chromatin accessibility during amphioxus development at single-cell resolution. Cell Reports 39, 110979, doi:10.1016/j.celrep.2022.110979 (2022).

54 Wang, W. et al. A single-cell transcriptional roadmap for cardiopharyngeal fate diversification. Nat Cell Biol 21, 674–686, doi:10.1038/s41556-019-0336-z (2019).

55 Sharma, S., Wang, W. & Stolfi, A. Single-cell transcriptome profiling of the Ciona larval brain. Dev Biol 448, 226–236, doi:10.1016/j.ydbio.2018.09.023 (2019).

56 Erkenbrack, E. M. et al. Whole mount in situ hybridization techniques for analysis of the spatial distribution of mRNAs in sea urchin embryos and early larvae. Methods Cell Biol 151, 177–196, doi:10.1016/bs.mcb.2019.01.003 (2019).

57 Hasegawa, Y., Mark Welch, J. L., Rossetti, B. J. & Borisy, G. G. Preservation of three-dimensional spatial structure in the gut microbiome. PLoS One 12, e0188257, doi:10.1371/journal.pone.0188257 (2017).

58 Redmayne, N. & Chavez, S. L. Optimizing Tissue Preservation for High-Resolution Confocal Imaging of Single-Molecule RNA-FISH. Curr Protoc Mol Biol 129, e107, doi:10.1002/cpmb.107 (2019).

59 Wei, J. et al. EDomics: a comprehensive and comparative multi-omics database for animal evo-devo. Nucleic Acids Res 51, D913–d923, doi:10.1093/nar/gkac944 (2023).

60 Fritzsch, B. & Straka, H. Evolution of vertebrate mechanosensory hair cells and inner ears: toward identifying stimuli that select mutation driven altered morphologies. Journal of Comparative Physiology A 200, 5–18, doi:10.1007/s00359-013-0865-z (2013).

61 Fritzsch, B. & Elliott, K. L. Gene, cell, and organ multiplication drives inner ear evolution. Dev Biol 431, 3–15, doi:10.1016/j.ydbio.2017.08.034 (2017).

62 Streit, A. Origin of the vertebrate inner ear: evolution and induction of the otic placode. J Anat 199, 99–103, doi:10.1046/j.1469-7580.2001.19910099.x (2001).

63 Lacalli, T. C. Sensory systems in amphioxus: a window on the ancestral chordate condition. Brain Behav Evol 64, 148–162, doi:10.1159/000079744 (2004).

64 Bone, Q. & Ryan, K. P. Cupular sense organs in Ciona (Tunicata: Ascidiacea). Journal of Zoology 186, 417–429, doi:10.1111/j.1469-7998.1978.tb03931.x (1978).

65 Mackie, G. O. & Singla, C. L. Cupular organs in two species of Corella (Tunicata: Ascidiacea). Invertebrate Biology 123, 269–281, doi:10.1111/j.1744-7410.2004.tb00161.x (2004).

66 Burighel, P. et al. Novel, secondary sensory cell organ in ascidians: In search of the ancestor of the vertebrate lateral line. Journal of Comparative Neurology 461, 236–249, doi:10.1002/cne.10666 (2003).

67 Mackie, G. O., Burighel, P., Caicci, F. & Manni, L. Innervation of ascidian siphons and their responses to stimulation. Canadian Journal of Zoology 84, 1146–1162, doi:10.1139/z06-106 (2006).

68 Gasparini, F. et al. Cytodifferentiation of hair cells during the development of a basal chordate. Hearing research 304, 188–199, doi:10.1016/j.heares.2013.07.006 (2013).

69 Dayaratne, M. W., Vlajkovic, S. M., Lipski, J. & Thorne, P. R. Kolliker’s organ and the development of spontaneous activity in the auditory system: implications for hearing dysfunction. Biomed Res Int 2014, 367939, doi:10.1155/2014/367939 (2014).

70 Delsuc, F., Brinkmann, H., Chourrout, D. & Philippe, H. Tunicates and not cephalochordates are the closest living relatives of vertebrates. Nature 439, 965–968, doi:10.1038/nature04336 (2006).

71 Fredriksson, G., Öfverholm, T. & Ericson, L. E. Electron-microscopic studies of iodine-binding and peroxidase activity in the endostyle of the larval amphioxus (*Branchiostoma lanceolatum*). Cell and Tissue Research 241, 257–266, doi:10.1007/BF00217169 (1985).

72 Onuma, T. A., Nakanishi, R., Sasakura, Y. & Ogasawara, M. Nkx2-1 and FoxE regionalize glandular (mucus-producing) and thyroid-equivalent traits in the endostyle of the chordate *Oikopleura dioica*. Developmental Biology 477, 219–231, doi:10.1016/j.ydbio.2021.05.021 (2021).

73 Yamagishi, M. et al. Differentiation of endostyle cells by Nkx2-1 and FoxE in the ascidian *Ciona intestinalis* type A: insights into shared gene regulation in glandular- and thyroid-equivalent elements of the chordate endostyle. Cell and Tissue Research 390, 189–205, doi:10.1007/s00441-022-03679-w (2022).

74 Ozaki, T. et al. Thyroid regeneration: characterization of clear cells after partial thyroidectomy. Endocrinology 153, 2514–2525, doi:10.1210/en.2011-1365 (2012).

75 Hao, Y. et al. Integrated analysis of multimodal single-cell data. Cell 184, 3573–3587 e3529, doi:10.1016/j.cell.2021.04.048 (2021).

76 McGinnis, C. S., Murrow, L. M. & Gartner, Z. J. DoubletFinder: Doublet Detection in Single-Cell RNA Sequencing Data Using Artificial Nearest Neighbors. Cell Systems 8, 329–337.e324, doi:10.1016/j.cels.2019.03.003 (2019).

77 Young, M. D. & Behjati, S. SoupX removes ambient RNA contamination from droplet-based single-cell RNA sequencing data. Gigascience 9, doi:10.1093/gigascience/giaa151 (2020).

78 Stirling, D. R. et al. CellProfiler 4: improvements in speed, utility and usability. BMC Bioinformatics 22, 433, doi:10.1186/s12859-021-04344-9 (2021).

79 Tarashansky, A. J. et al. Mapping single-cell atlases throughout Metazoa unravels cell type evolution. Elife 10, doi:10.7554/eLife.66747 (2021).

80 Cao, J. et al. The single-cell transcriptional landscape of mammalian organogenesis. Nature 566, 496–502, doi:10.1038/s41586-019-0969-x (2019).

81 Qiu, X. et al. Mapping transcriptomic vector fields of single cells. Cell 185, 690–711 e645, doi:10.1016/j.cell.2021.12.045 (2022).

82 Jin, S. et al. Inference and analysis of cell-cell communication using CellChat. Nat Commun 12, 1088, doi:10.1038/s41467-021-21246-9 (2021).

83 Cosentino, S. & Iwasaki, W. SonicParanoid: fast, accurate and easy orthology inference. Bioinformatics 35, 149–151, doi:10.1093/bioinformatics/bty631 (2019).

84 Armingol, E. et al. Inferring a spatial code of cell-cell interactions across a whole animal body. PLoS Comput Biol 18, e1010715, doi:10.1371/journal.pcbi.1010715 (2022).

85 Qiu, X. et al. Single-cell mRNA quantification and differential analysis with Census. Nat Methods 14, 309–315, doi:10.1038/nmeth.4150 (2017).

